# Encephalitic alphavirus infection reduces neurite outgrowth and downregulates Rho GTPase pathways in human iPSC-derived neurospheres

**DOI:** 10.64898/2026.10.08.755254

**Authors:** K Cirksena, LM Haiber, BA Kühne, J Hupfeld, M Duven, A Fischer, M Schenke, F Richter, S Weiß, M Altenbuchinger, N Seifert, HU Zacharias, S Konzok, B Seeger, G Gerold

**Affiliations:** Department of Biochemistry & Research Center for Emerging Infections and Zoonoses (RIZ), University of Veterinary Medicine Hannover, Hannover, Germany; Department of Cellular Proteome Research, Helmholtz Centre for Infection Research (HZI), Braunschweig, Germany; Institute for Food Quality and Food Safety, Research Group Food Toxicology and Alternative/Complementary Methods to Animal Experiments, University of Veterinary Medicine Hannover, Hannover, Germany; Department of Pharmacology, Toxicology, and Pharmacy, University of Veterinary Medicine Hannover, Hannover, Germany; German Federal Institute for Risk Assessment (BfR), German Centre for the Protection of Laboratory Animals (Bf3R), Berlin, Germany; Institute of Virology, Medical School Hannover, Hannover, Germany; Institute of Pharmacology and Toxicology, School of Veterinary Medicine, Freie Universität Berlin, Berlin, Germany; School of Science, Constructor University, Bremen, Germany; Center for Systems Neuroscience (ZSN), Hannover, Germany; Department of Medical Bioinformatics, University Medical Center Göttingen, Göttingen, Germany; Peter L. Reichertz Institute for Medical Informatics of TU Braunschweig and Hannover Medical School, Hannover, Germany; Fraunhofer Institute for Toxicology and Experimental Medicine, Preclinical Pharmacology and Toxicology, Nikolai-Fuchs-Str. 1, Hannover, Germany; Department of Clinical Microbiology, Umeå University, Umeå, Sweden; Wallenberg Centre for Molecular Medicine (WCMM), Umeå University, Umeå, Sweden; Department of Hygiene, Microbiology and Virology, Institute of Virology, Medical University of Innsbruck, Innsbruck, Austria

## Abstract

Venezuelan equine encephalitis virus (VEEV) is a re-emerging neurotropic arbovirus, which affects the developing human brain, potentially leading to miscarriage, stillbirth, or long-term neurological sequelae. Currently, the mechanisms underlying these sequelae remain largely elusive. Using human induced pluripotent stem cell (iPSC)-derived neurospheres as a model for brain development, we show that both human neurons and glial cells are susceptible to VEEV infection. Microscopy-based quantification reveals that VEEV impairs neurite outgrowth in the human neurospheres. To mechanistically assess these observations, we performed multi-omics analyses of VEEV-infected neurospheres and demonstrate translational shutoff including downregulation of Rho GTPase pathways. The transcriptional and proteome profile is indicative of reduced actin cytoskeleton dynamics and consequently migratory defects of cells in the neurospheres. Shotgun secretomics moreover revealed downmodulation of neurite outgrowth associated proteins including midkine (MDK), amyloid-beta precursor protein (APP) and microtubule-associated protein 1B (MAP1B). Our findings reveal mechanistic insights into how VEEV infection impairs neuronal development and demonstrate that neurospheres are a powerful platform for assessing teratogenic effects of viruses on brain development.

## Introduction

Venezuelan equine encephalitis virus (VEEV) is a neurotropic, mosquito-borne alphavirus, which infects the central nervous system (CNS) and is highly pathogenic. Endemic to the Americas, VEEV is a growing public health threat due to its dual role as a re-emerging zoonosis and a Category B biological agent. During natural infection, the enzootic transmission cycle involves rodents and mosquitoes, while equids act as amplification hosts during epizootics, facilitating spillover into human populations^1^. In humans, VEEV infection typically causes an acute febrile illness which, in 5-15% of cases, progresses to encephalitis with marked neuronal death, neuroinflammation, and disruption of the blood–brain barrier^1, 2^. Although overall human mortality is low (< 1%), long-term neurological sequelae — including seizures, cognitive deficits, motor and sensory impairments — occur in 4–14% of survivors. Teratogenic effects often lead to stillborn deaths and children under the age of five years have the highest risk of neurological sequelae of up to 35%^1, 3^.

The age-dependent vulnerability to VEEV mirrors observations in animal models, where mid-gestational infection in rodents induces placental inflammation and near-complete fetal loss^4^, while non-human primate studies reveal microcephaly, hydrocephalus, and cataracts in surviving offspring^5^. Human case reports further corroborate developmental risks, documenting VEEV-positive fetal brain tissue, spontaneous abortions, and neonates with microcephaly or cortical necrosis^6, 7, 8^. Collectively, these findings underscore the developing brain’s unique susceptibility to VEEV — a critical yet mechanistically incompletely resolved aspect of its pathogenesis.

Current insights into VEEV neuropathogenesis stem predominantly from rodent *in vivo* models^9, 10, 11^ or from reductionist *in vitro* systems using tumor-derived human cell lines or primary CNS cells^10, 12^. The latter models have described neuronal and glial tropism, host translational shutoff, and pro-inflammatory cytokine secretion. However, they cannot fully account for the cellular diversity, intercellular signaling, and maturational trajectories inherent to human neurodevelopment. Notably, we lack studies leveraging human iPSC-derived models to investigate how VEEV infection disrupts fetal brain maturation — a gap that obscures its teratogenic mechanisms and hinders therapeutic development.

To address this gap, we established a human iPSC-derived 3D neurosphere model. This model comprises neural progenitor cells, neuronal and glial cells, and recapitulates key CNS developmental processes, within an architecturally complex, physiologically relevant microenvironment^13^. Using a GFP reporter virus, we showed that VEEV infects both human neurons and glial cells. VEEV infection impaired neurite outgrowth in human neurospheres, as assessed using a microscopic assay of plated spheres. In accordance with this phenotype, transcriptomic and proteomic analyses confirmed translational shutoff and identified downregulation of Rho GTPase pathways. We next established a secretomics workflow for infected neurospheres and unraveled that VEEV decreases secretion of the neurite outgrowth associated proteins MDK, APP and MAP1B. Together, our data help to address a critical gap in alphavirus teratogenicity research and provide a relevant model for studying pathogens affecting human neurodevelopment.

## Results

### Human iPSC-derived neurospheres display hallmarks of brain development

To develop a model for VEEV-dependent modulation of neural development, we generated iPSC-derived neurospheres and assessed their differentiation status. Stem cell-derived neuronal 3D models may contain various subsets of neural cell types, which depend on applied differentiation protocols. Immunofluorescence microscopy revealed that neurospheres contain both neuronal, tubulin-III-beta (TUJ1) and glial, glial fibrillary acid protein (GFAP)-positive cells (Fig. 1A). Further, we confirmed expression of cell type-specific markers involved in brain development via immunostaining. Specifically, neurospheres expressed SRY-Box Transcription Factor 2 (SOX2), nestin, doublecortin (DCX), microtubule-associated protein 2 (MAP2), and S100 calcium-binding protein B (S100B) (Supplementary Fig. 1A-C). For in depth characterization of cellular markers, we performed transcriptome and whole cell proteome analysis of floating neurospheres. In total, we identified 36,612 transcripts in 4 measured samples and 5,141 proteins in 8 samples. We initially compared the neurosphere proteome to that of iPSCs with 4,397 proteins in 4 replicates. We observed a strong decrease of the pluripotency marker protein lin-28 homolog A (LIN28) upon iPSC differentiation to neurospheres and a dominant upregulation of proteins expressed during brain development (Supplementary Fig.1D). In neurospheres, we further identified markers^13^ for neural progenitor cells as well as neurons and glial cells (Fig. 1B). We found evidence for both immature and mature neurons, the latter including cholinergic, dopaminergic, GABAergic, glutamatergic, glycinergic, noradrenergic and serotonergic neurons (for comparison see Lislien et al. (2025)^14^). Additionally, selected genes expressed during brain development were upregulated on mRNA level during differentiation from iPSCs to neurospheres (Supplementary Fig. 1E).

**Fig. 1:**
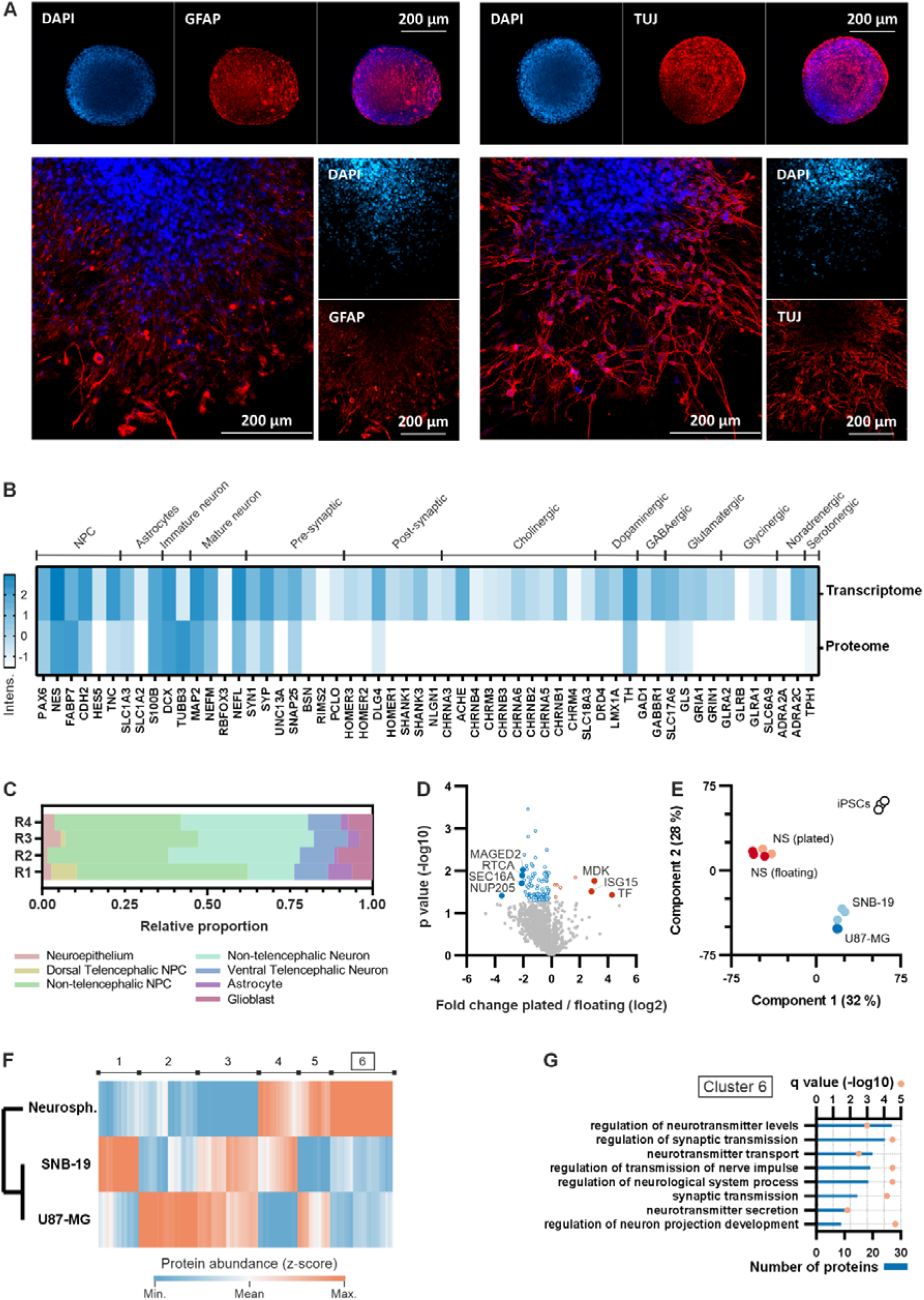
Floating and plated neurospheres display brain development-related gene and protein expression profiles distinct from pluripotent stem cells and glioblastoma cell lines. **(A)** Confocal immunofluorescence microscopy of floating (upper panel) and plated neurospheres (lower panel) stained for GFAP, TUJ1 and DNA (DAPI). Scale bar: 200 µm. **(B)** Selected markers for different neural cell types (indicated on top) expressed in floating neurospheres and identified via RNAseq or LC-MS analysis (n = 4). Z-transformed mean intensities of transcripts (fpkm ≥ 1) or proteins (iBAQ > 0) detected in at least one replicate are represented by blue colour (darker blue = higher expression, white = not detected). (**C**) Cell type-associated transcriptional patterns detectable in neurospheres derived from comparison of bulk RNAseq data (fpkm) with the transcriptomic cell atlas of human neural organoids (supervised deconvolution). IP = intermediate progenitor, NPC = neural progenitor cell, R = replicate. **(D)** Whole cell proteome of floating neurospheres cultured plated on ECM compared to that of floating neurospheres by pairwise t-tests (n = 3, 2,070 proteins). Proteins with significantly altered abundance in plated spheres (p ≤ 0.05, 98 proteins) are highlighted in blue (reduced) or red (increased), those with absolute fold changes ≥ 4 are labelled. **(E)** Principal component analysis (n = 3, 4,042 proteins) comparing the overall whole cell proteomes of stem cells (iPSCs, white dots), stem cell-derived neurospheres (NS, cultured floating or plated, red dots) and two glioblastoma cell lines (SNB-19 and U87-MG, blue dots). **(F-G)** Whole cell proteome analysis (n = 4, 3,545 proteins) comparing floating neurospheres and glioblastoma cells (SNB-19, U87-MG). Differentially expressed proteins were defined by one-way ANOVA (FDR ≤ 0.05). (F) Abundance of differentially expressed proteins displayed by z-transformed mean LFQ intensities after hierarchical clustering (2,902 proteins). The six visible main clusters (labelled on top) comprise proteins with increased (red) or decreased (blue) expression in one of the three cellular models. (G) Protein functions (GOBP) enriched among proteins of cluster 6 as shown in (F) (619 proteins with significantly higher abundance in neurospheres compared to glioblastoma cells). Bars display the number of proteins with the respective protein function. Q values are displayed by red dots. (Fisher exact test, q ≤ 0.05, relative to all differentially expressed proteins, most significant (by q) results only).

To further characterize the cellular composition of neurospheres, we performed supervised deconvolution of the bulk RNAseq data using a reference single-cell transcriptomic atlas of human neural organoids^15^. We estimated the contribution of cell-type–associated transcriptional programs by comparing bulk expression profiles to selected reference cell states. Across replicates, neural progenitor– and neuron-associated signatures dominated, with additional contributions from astrocyte-like programs (Fig. 1C). In sum, these results confirm that our neurospheres recapitulate key features of a developing neural system.

While floating neurospheres display 3D architecture, specific biological functions such as cell migration require plating of neurospheres on an extracellular matrix (ECM)-coated surface. Hence, we next compared the proteomes of floating neurospheres (floating) with that of neurospheres plated on ECM (plated, 3,702 proteins, 3 replicates). We identified seven differentially expressed proteins with absolute fold changes ≥ 4 (p ≤ 0.05). Among these, the neuronal migration promoting protein midkine (MDK)^16^ showed increased expression in plated neurospheres while the axon regeneration inhibitor RNA 3’-terminal phosphate cyclase (RTCA) showed decreased expression^17^ (Fig. 1D). Finally, we compared the neurosphere model to two CNS-derived cell lines commonly used in studies with neurotropic viruses, namely the glioblastoma cells lines SNB-19 and U87-MG (4,488 and 4,586 proteins, respectively, 4 samples each). The iPSC proteome served as additional reference here. Protein expression patterns were generally comparable among floating and plated neurospheres but clearly deviated from those of glioblastoma cells or iPSCs as determined by principal component analysis (Fig. 1E). In fact, among the 3,545 proteins compared between neurospheres and glioblastoma cells, 2,902 proteins were differentially expressed (ANOVA, FDR ≤ 0.05). Hierarchical clustering of these proteins revealed six distinct clusters with cell model-specific expression patterns (Fig. 1F) and downstream enrichment analysis revealed differences in fundamental biological pathways (Supplementary Fig. 1F-G). Notably, neurospheres distinctively expressed proteins involved in synaptic processes (Fig. 1G). Taken together, neurospheres contain neural progenitor cells, immature and mature neuronal and glial populations and express cell-type specific markers characteristic of the developing human brain.

### VEEV infects glial cells and neurons in iPSC-derived neurospheres

Next, we sought to evaluate the potential of iPSC-derived neurospheres as a model to study the effect of neurotropic viruses on the developing brain. To that end, we performed time-resolved infection experiments with VEEV encoding a GFP reporter and analyzed cell tropism, viral replication and host proteome changes. We observed that VEEV infected TUJ1 and GFAP-positive cells, both in neurospheres plated on ECM as well as in floating spheres (Supplementary Fig. 2A). Infectious doses ranged from 3 to 300 FFU per sphere for floating spheres (Supplementary Fig. 2A) and 5 to 450 FFU per sphere for plated spheres (Supplementary Fig. 2B). Exemplary GFP signals in floating neurospheres 24 hours post infection are shown for conditions with and without serum supplementation (Supplementary Fig. 2C). To get first insights into virus-host interactions in floating neurospheres, we infected neurospheres with VEEV (30 FFU) followed by sampling between 6 and 48 hours post infection (hpi) with spheres harvested prior to infection (0 hpi) as a control (Supplementary Fig. 2D). We quantified the abundance of intracellular viral genomes and proteins by qPCR and LC-MS, respectively. Viral replication was consistent across all analyzed samples with first genome copies detected at 6 hpi and genome copies reaching a plateau at 24 hpi (Fig. 2B). In contrast, we identified intracellular viral structural and nonstructural proteins starting at 24 hpi and accumulating up to 48 hpi (Fig. 2C). We next assessed if VEEV not only replicates its RNA genome in human neurospheres but also assembles new infectious progeny particles and releases these into culture supernatants. To that end, we washed neurosphere cultures extensively after infection and sampled supernatants at different timepoints post infection. These supernatants were then titrated on susceptible SNB-19 cells. We detected few secreted infectious particles already at 10 hpi (2 FFU per neurosphere) and infectious titers in supernatants increased up to 48 hpi to 1.3 × 10^4^ FFU per neurosphere (Fig. 2D). Thus, VEEV completes the full viral life cycle in neurospheres.

**Fig. 2:**
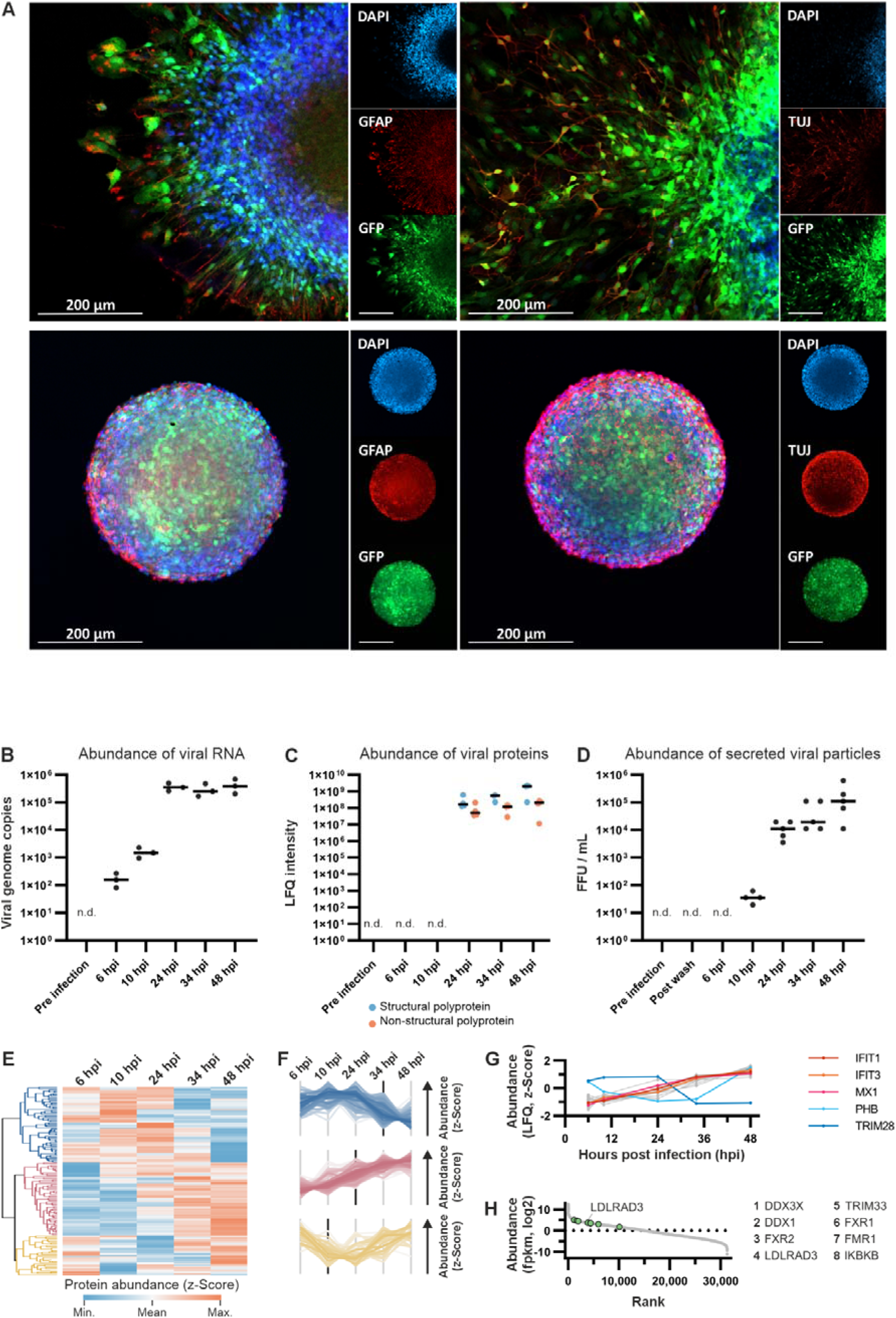
VEEV infects floating and plated neurospheres. **(A)** Confocal immunofluorescence microscopy of VEEV-infected (24 hpi; GFP) floating (lower panel, 300 FFU/sphere) and plated spheres (upper panel, 450 FFU/sphere) stained for GFAP (left panel), TUJ1 (right panel) and DNA (DAPI). Scale bar: 200 µm. **(B-D)** Viral replication post infection in floating neurospheres (30 FFU/sphere). n.d. = not detected. (B) Number of viral genome copies detected by qPCR at different time points post infection. Values were normalized to total RNA content per sample. Dashes represent the median of three biological replicates. (C) Abundance of viral proteins (LFQ) detected by LC-MS at indicated time points post infection. Dashes represent the median of four biological replicates. (D) Number of secreted infectious viral particles detected via TCID_50_ assay at different time points post infection. Dashes represent the FFU median of five biological replicates. **(E-G)** Whole cell proteome analysis of floating neurospheres 6-48 h post VEEV infection (30 FFU/sphere) analyzed by LC-MS (n = 4, 2,678 proteins). Differentially expressed proteins were determined by one-way ANOVA (p ≤ 0.05, 155 proteins). All graphs show z-transformed LFQ intensities. (E) Heat map showing abundance of differentially expressed proteins over time. Three main clusters of proteins are highlighted in blue, red and yellow. (F) Expression profiles of the proteins belonging to the clusters highlighted in (E). The proximity of a single protein’s abundance to the mean abundance of all cluster proteins is reflected by the intensity of the colour. Black dashes indicate the time post infection at which the major change of the respective protein expression profile becomes visible. (G) Expression profiles of differentially expressed proteins involved in defence mechanisms against viruses (Gene Ontology GO:0009615, selected proteins highlighted in red and blue, median). **(H)** Ranked abundance of all transcripts detected in uninfected floating neurospheres by RNAseq analysis (fpkm intensities, highest intensity = rank 1, 31,301 transcripts). Transcripts, known to be required for VEEV infection of cells are highlighted in green, numbers in the legend indicate the ranked intensity of the respective transcript (1 = highest intensity). The VEEV receptor LDLRAD3 is labelled.

With regards to VEEV-induced alterations of the host proteome, we observed major changes between 10 and 34 hpi (2,678 cellular proteins, Fig. 2E). Proteins clustered by three phenotypes: (I) proteins with decreased abundance upon VEEV infection, proteins with increased abundance upon infection, and proteins with early decrease and late recovery of expression (Fig. 2F). VEEV induced expression of selected proteins involved in antiviral defense (Fig. 2G). Expectedly, we detected transcripts of known VEEV host factors, i.e. transcripts for proteins required by the virus to complete its life cycle in neurospheres. These included the receptor low-density lipoprotein receptor class A domain-containing protein 3 (LDLRAD3) (Fig. 2H). Protein abundance of identified host factors did not change during infection (Supplementary Fig. 2E). Of note, there was no evidence for apoptotic processes induced by VEEV infection (Supplementary Fig. 2F). In sum, VEEV infected neuronal and glial cell populations and completed its full replication cycle in this multicellular neural model.

### VEEV reduces neurite outgrowth

Given that neurological sequelae following VEEV infection may result from impaired neuronal network formation, we hypothesized that VEEV could interfere with neurite outgrowth. Hence, we set out to develop a neurite outgrowth assay that assessed the length of neurites growing out of plated spheres (Fig. 3A). Such plated neurosphere assays are typically used to assess developmental neurotoxicity upon chemical exposure^18, 19, 20^. We adapted this assay to investigate effects of virus infection on the developing nervous system. To ensure that experiments were not confounded by cytotoxicity, we first evaluated cell viability and cytotoxicity upon infection. At 24 hpi, none of the tested infectious doses (5, 45 or 450 FFU per sphere) reduced viability or induced cytotoxicity (Fig. 3B). In contrast, at 48 hpi, viability was significantly reduced upon infection with 45 and 450 FFU per sphere (p ≤ 0.05, vs. uninfected controls). Notably, even the lowest infectious dose (5 FFU per sphere) exhibited reduced viability (71%) at 48 hpi despite weak GFP signal upon infection. Based on these findings, we chose the 24 hpi time point for subsequent experiments. This time point should primarily reflect early virus-induced mechanisms rather than secondary effects of metabolic collapse or pre-lethal stress.

**Fig. 3:**
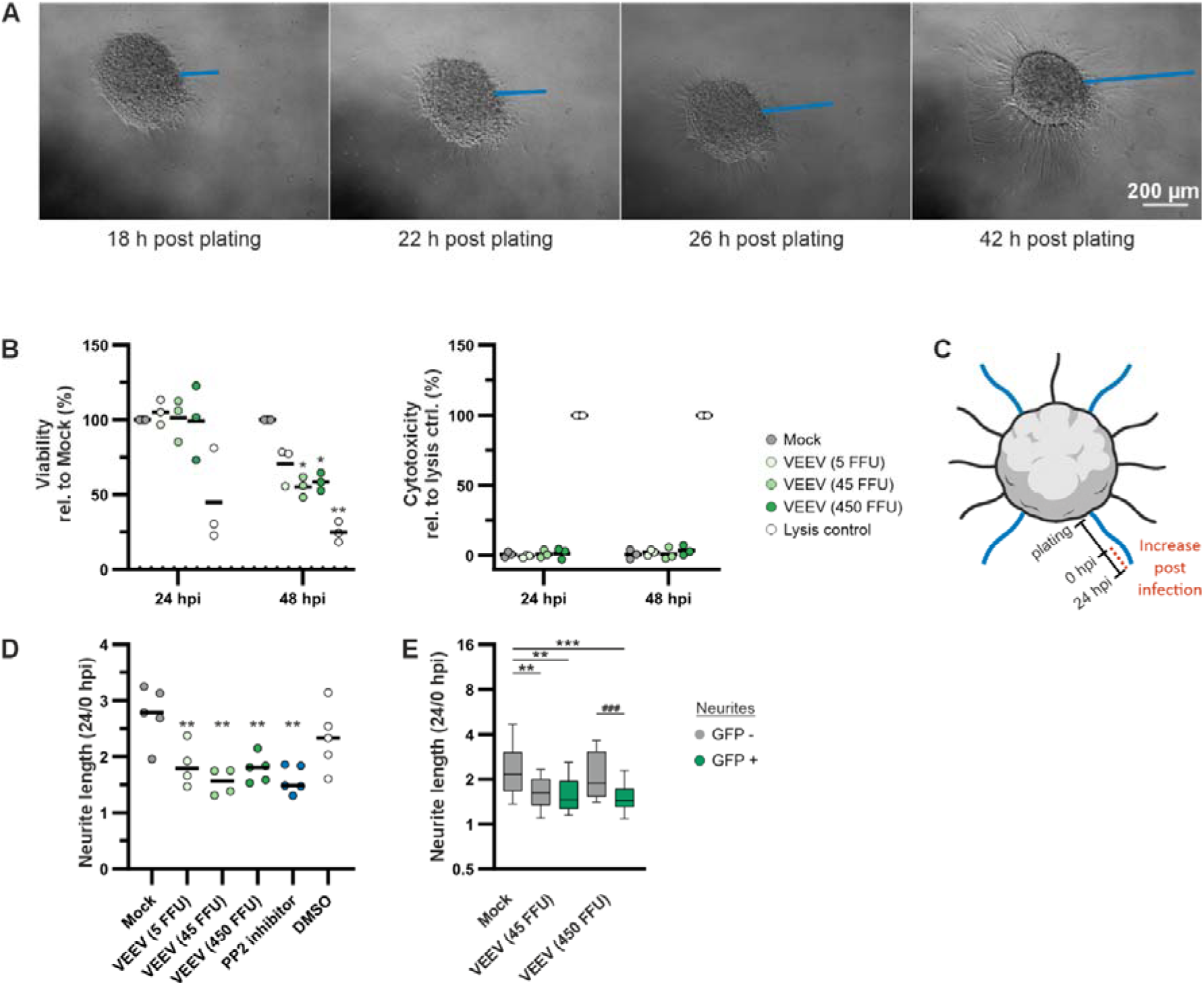
VEEV infection of neurospheres reduces neurite outgrowth. Single plated neurospheres were infected with 3 infectious doses of VEEV (5, 45 and 450 FFU). A migration inhibitor (PP2), DMSO and uninfected cells (Mock) served as controls. **(A)** Example of neurites growing out of plated neurospheres. The length of the same neurite (blue line) was measured at different time points to quantify neurite outgrowth. **(B)** Viability of neurospheres (left panel) and VEEV-induced cytotoxicity (right panel) upon infection determined by CTB and LDH assays, respectively. Recorded intensities were normalized to uninfected condition (viability) or lysis controls (cytotoxicity) after background subtraction. Dashes represent means calculated based on three separate experiments with 3-5 technical replicates. Means of different conditions were compared to 100% viability by one-sample T-tests (viability) or one-way ANOVA (cytotoxicity). **(C)** Scheme of migration assay. The length of four neurites was determined before infection (0 hpi) and 24 h post infection (24 hpi) to measure the increase of neurite lengths influenced by VEEV infection. Assay was performed 24 h post plating. **(D)** Neurite outgrowth of untreated spheres (Mock) compared to VEEV-infected, migration inhibitor-treated (PP2) or DMSO-treated (inhibitor solvent) spheres. Neurite lengths measured 24 h post treatment were normalized to neurite lengths measured before treatment. Dashes show the means of 5 individual experiments, in which with 3-5 spheres per condition and 4 neurites per sphere were averaged (12-52 measured neurites per condition and experiment as shown in Fig. S3C). Each condition was compared to Mock condition by one-way ANOVA. **(E)** Neurite outgrowth, as determined in (D). Neurites of selected GFP-positive sites were compared to selected GFP-negative sites within infected spheres by pairwise T-tests (20-67 measured neurites per group, significance indicated by asterisks). Neurites of infected spheres were compared to Mock-derived neurites by one-way ANOVA (significance indicated by hashes). Boxes represent the 25th–75th percentiles, the center line indicates the median (50th percentile), and whiskers represent the 10th–90th percentiles. (Statistics) Asterisks and hashes indicate significance levels of 0.05 (*/#), 0.005 (**/##), 0.0005 (***/###).

Next, we determined neurite lengths at four different loci of a neurosphere 24 hours post plating before (0 hpi) and after (24 hpi) VEEV infection in five independent experiments (Fig. 3C, 796 neurites in total). The length of neurites across untreated spheres determined 48 h post plating (corresponds to 24 hpi) varied between 36 and 667 µm. We recognized clear differences in length both within the same sphere as well as between different spheres and batches (Supplementary Fig. 3A). To account for this variation, we first established a reproducible data analysis method using the cell migration inhibitor PP2 targeting Src kinase. While the effect of migration inhibitor treatment was barely visible by comparing the raw neurite lengths determined 0 hpi or 24 hpi, the comparison of neurite lengths measured 24 hpi normalized to lengths measured 0 hpi, i.e. the quantification of neurite outgrowth, resulted in significant differences between untreated spheres and spheres treated with the Src kinase inhibitor (Supplementary Fig. 3B). Upon inhibitor treatment, neurite outgrowth was reduced to 58%. We then studied the influence of VEEV infection on neurite outgrowth by infecting spheres with 5-450 FFU per neurosphere. Compared to untreated spheres, neurite outgrowth in infected spheres was reduced in a dose dependent manner for low to intermediate infectious doses and up to 56% (Fig. 3D, Supplementary Fig. 3C). Of note, VEEV infection and the PP2 inhibitor reduced neurite outgrowth to similar degree. Finally, as VEEV only infected a fraction of neurons in each sphere, we asked if specifically, the VEEV infected, GFP positive neurons were impaired in neurite outgrowth or also bystander, GFP negative neurons. At the highest infection dose (450 FFU), neurite outgrowth impairment was more pronounced in GFP positive neurons compared to GFP negative neurons. However, at the lower infection dose (45 FFU), neurite outgrowth was reduced in GFP-positive and GFP-negative neurons in a comparable manner (Fig. 3E). Thus, VEEV reduced neurite outgrowth in infected neurons and -to a lesser degree- in bystander neurons and this effect was independent of viral cytopathic effects.

### VEEV infection of neurospheres leads to translational shutoff and downmodulates Rho GTPase pathways

To unravel expression pattern changes induced by VEEV infection that may lead to impairment of neurite outgrowth, we performed an extensive multi-omics experimental series. We chose the 24 hpi timepoint at which we had previously excluded a cytopathic effect. Next, we compared VEEV-infected neurospheres, to non-infected controls on transcriptome and proteome level (39,596 identified transcripts, 6,734 proteins, 4 replicates per condition). Viral RNA genomes were detected in all infected samples. However, the number of identified viral RNA genomes and the measured abundance of viral proteins varied considerably between biological replicates (Supplementary Fig. 4A-B). VEEV induced changes at both transcriptome and cellular proteome level with moderate differences between samples derived from separate sets of differentiated neurospheres (Supplementary Fig. 4C).

While VEEV infection predominantly upregulated host transcript levels, it led to more balanced up- and downregulation on protein level with overall less pronounced effects (Fig. 4A-B). In the transcriptome, 725 out of 981 (74%) differentially expressed transcripts showed increased expression, while in the proteome, 107 out of 201 (53%) significantly affected proteins exhibited increased abundance (both p ≤ 0.05). VEEV infection upregulated the interferon-induced proteins ubiquitin-like protein ISG15 (ISG15), interferon-induced protein with tetratricopeptide repeats 3 (IFIT3) and interferon-induced 35 kDa protein (IFI35) on RNA and protein level (Fig. 4A-B). In contrast, VEEV downmodulated ATP-binding cassette sub-family D member 3 (ABCD3), involved in lipid metabolism and discussed as an effector of Rho GTPases^21^ on transcript and protein level (Fig. 4A-B). Pathway analyses highlighted immune activation at RNA level in contrast to proteostasis and cell signaling changes at protein level (Supplementary Fig. 4D-E).

**Fig. 4:**
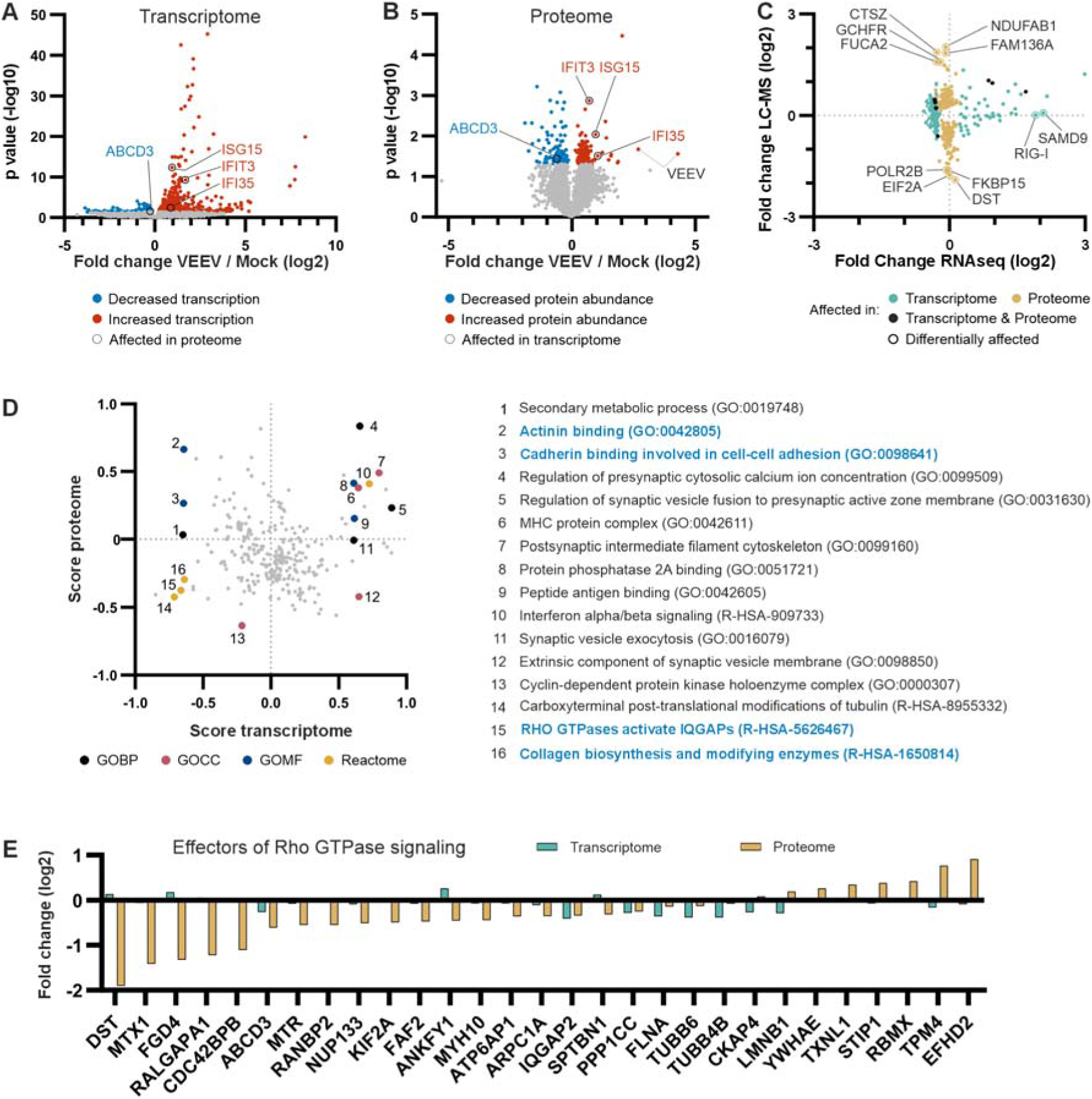
VEEV induces a translational shutoff and downregulates RhoGTPases in floating neurospheres. Transcriptome and whole cell proteome analyses of neurospheres upon VEEV infection (24 hpi, 30 FFU/sphere, n = 4). Differentially expressed transcripts and proteins were defined by pairwise t-tests (p ≤ 0.05). (**A-B**) Volcano plots showing overall transcriptome (A, 35,543 transcripts) and whole cell proteome (B, 2,894 proteins) changes in neurospheres upon infection compared to uninfected cells. Differentially expressed transcripts (981) and proteins (201) are highlighted in blue (reduced expression) and red (increased expression). Genes significantly and equally affected on both levels of gene expression are labelled and highlighted by circles. Identified VEEV proteins (structural and non-structural) are labelled in black. **(C)** VEEV-induced fold changes of 277 differentially expressed genes quantified on both, transcript and protein level. Yellow: significantly altered transcription; turquoise: significantly altered protein abundance; black: differentially expressed on transcript and protein level. Genes differentially affected on transcript and protein level (minimum 1.5-fold change one level and maximum 0.5-fold change on the other) are labelled and highlighted by circles. **(D)** 2D enrichment analysis performed with VEEV-induced fold changes (2860 transcripts / proteins) detectable by both RNAseq and LC-MS. Results were filtered for cellular processes whose members received significantly (p ≤ 0.05 and FDR ≤ 0.4) higher or lower fold changes compared to the average of all observed fold changes. The score reflects the average fold change observed for the respective cellular process. The most 4 significant processes (by p value, Score > 0.6) of databases GOBP Slim, GOCC slim, GOMF and Reactome are coloured and labelled by numbers with their respective names listed right of the plot. Processes potentially related to neural cell migration or neurite outgrowth are highlighted in blue. **(E)** Fold changes of all proteins annotated as Rho GTPase signaling effectors (child IDs of R-HSA-194315 in Reactome database) identified in neurospheres by both RNAseq (turquoise) and LC-MS (yellow) analysis and differentially expressed on RNA or protein level. Bars represent medians.

To disentangle overlapping gene-expression programs in the bulk transcriptome and assess whether infection-induced changes could be influenced by cellular composition, we deconvoluted the RNAseq data. Using unsupervised deconvolution we grouped co-regulated genes by shared expression patterns across all infected replicates. We identified four gene-expression programs and a reproducible redistribution upon VEEV infection (Supplementary Fig. 4F). VEEV infection induced an antiviral/immune-associated program and reduced protein synthesis (translation) processes (Supplementary Fig. 4G). Notably, we did not find apoptosis-associated transcriptional signatures at transcriptomic level (Supplementary Fig. 4H). VEEV-induced changes were not attributable to gene-expression patterns specific to individual cell populations, except for only limited overlap with neural progenitor-associated gene expression (Supplementary Fig. 4I). Supervised deconvolution based on a reference single cell dataset revealed only modest shifts in cell type-associated signatures upon infection, with no evidence for decreased neuronal or glial signatures (Supplementary Fig. 4J). Together, these data indicate that VEEV induces coordinated changes in gene-expression programs rather than substantial loss of specific cell populations.

Alphaviruses are known to induce a translational shutoff^22^. In line with this, the comparison of fold changes of significantly altered transcripts and proteins revealed that VEEV-induced effects on transcript and protein level were decoupled (Fig. 4C). For instance, the antiviral genes sterile alpha motif domain-containing protein 9 (SAMD9) and the RNA sensor retinoic acid-inducible gene 1 protein (DDX58/RIG-I) increased on transcript but remained unchanged on protein level. This translational shutoff mainly affected antiviral interferon stimulated genes (ISGs), while VEEV proteins were, as expected, expressed at high levels (Fig. 4B). In addition, we identified four genes (FK506-binding protein 15 (FKBP15), dystonin (DST), eukaryotic translation initiation factor 2A (eIF2A) and POLR2B), for which VEEV did not affect transcript levels but decreased protein levels. FKBP15 and DST are part of the cytoskeleton and cellular trafficking machinery with FKBP15 (or FKBP133) being important for the cytoskeletal organization of neuronal growth cones^23^. POLR2B and eIF2A are essential for initiation of transcription and translation, respectively^24, 25^. Of note, the 165 most affected transcripts (by absolute fold change) were not affected on protein level. VEEV infection also increased abundance of some proteins without altering respective transcript levels at the 24 hpi time point. These upregulated proteins are involved in mitochondrial processes, redox or lipid metabolism (acyl carrier protein, mitochondrial (NDUFAB1), protein FAM136A (FAM136A), GTP cyclohydrolase 1 feedback regulatory protein (GCHFR)).

Next, we compared VEEV-induced changes in neurospheres with those observed in the two glioblastoma cell lines, SNB-19 and U87-MG (Supplementary Fig. 4K-L). The cellular response to VEEV infection was highly specific to each experimental model, reflected by the low overlap of significantly altered proteins (Supplementary Fig. 4M-P). Nonetheless, some biological processes were altered in all three models, i.e. antiviral and stress responses and RNA metabolism regulation (Supplementary Fig. 4Q-S). It is noteworthy that VEEV altered retrograde transport from endosomes to the Golgi network only in neurospheres. VEEV affected six proteins in all tested models among them the two VEEV host factors FXR1 (FXR1)^26, 27^ and Ras GTPase-activating protein-binding protein 1 (G3BP1)^28, 29^. Interestingly, G3BP1 was downregulated in glioblastoma cells, but upregulated in neurospheres.

Finally, we performed a two-dimensional enrichment analysis to identify cellular processes in neurospheres regulated by VEEV on transcript and/or protein level (Fig. 4D). Strong induction on mRNA and protein level was visible for antiviral defense effectors (IFN signaling) as well as genes involved in various synaptic processes. In addition, we observed an overall reduction of factors of the cytoskeleton and Rho GTPase effectors, known as modulators of cell shape, motility and neurite outgrowth (Fig. 4D-E). Our findings demonstrate a VEEV-induced uncoupling of transcriptome and proteome in neurospheres. In sum, VEEV not only downmodulates protein abundance of a set of ISGs, the RNA polymerase II and the translation initiation factor eIF2A, but also of proteins that are involved in cytoskeletal functions needed for neurite outgrowth.

### VEEV infection of neurospheres reduces secretion of cognitive developmental factors

Given that VEEV affected neurite outgrowth in infected and bystander neurons, we speculated that this phenotype was not only caused by cell-intrinsic effects on cytoskeletal pathways, but also by paracrine signaling. Therefore, we set out to determine VEEV induced changes in protein secretion of neurosphere cells using LC-MS-based secretome analyses. As the host cell origin of a secreted protein is elusive in a mixed glial cell and neuron model such as neurospheres, we also analyzed the secretome of isolated single glioblastoma cell lines, SNB-19 and U87-MG cells. We chose to analyze protein secretion 24 hpi post infection with VEEV, again to avoid cytopathic effects. Since serum proteins can mask identification of secreted proteins, we established serum-free culture conditions. To estimate the effect of serum-depletion, we first compared transcriptomic profiles of neurospheres cultured under serum-free and standard conditions with or without VEEV infection. Serum-depletion alone strongly impacted intracellular gene expression, including a downregulation of numerous virus defense mediators. However, the transcription changes induced by serum-depletion did not impair the antiviral response induced by VEEV infection (Supplementary Fig. 5A). Thus, we proceeded to collect, enrich and quantify proteins secreted from VEEV-infected neurospheres and cell lines during a 24 h period.

The number of secreted proteins identified in the supernatants of floating neurospheres varied between 139 and 508 proteins across the four replicates analyzed. Final data analysis was restricted to two replicates and 128 proteins quantified in all uninfected or infected samples (Fig. 5A, Supplementary Fig. 5B). Secretome analyses of glioblastoma cells was improved by higher cell densities and yielded sets of 33/192 (SNB-19) and 259/2,568 (U87-MG) proteins (low/high cell density, Fig. 5C, Supplementary Fig. 5E-F). Importantly, most proteins (89–94%) analyzed across all secretome experiments were classified as extracellular, confirming the absence of significant cellular debris contamination upon serum depletion (Supplementary Fig. 5G).

**Fig. 5:**
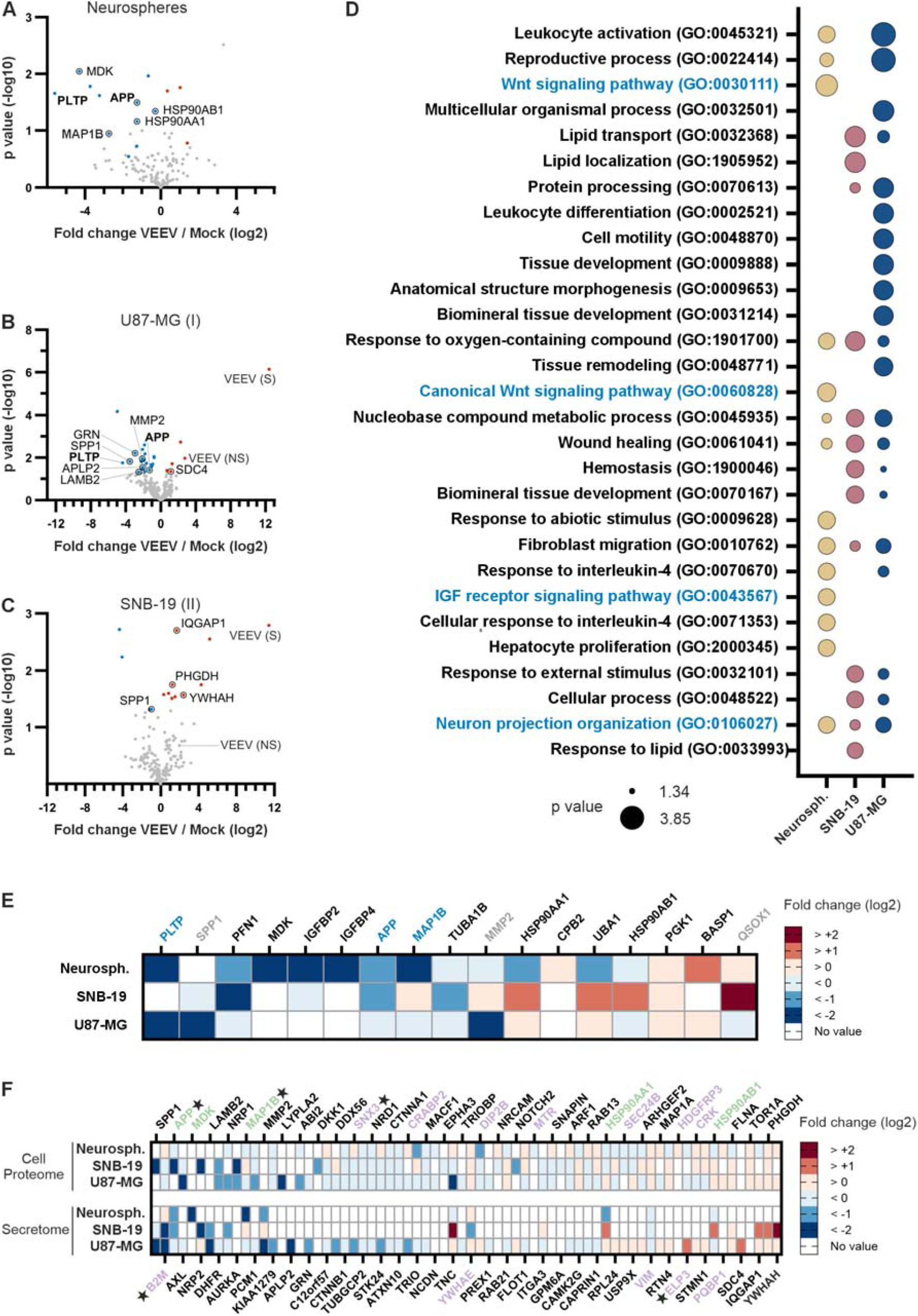
VEEV reduces secretion of cognitive development factors. Secretome analysis of VEEV-infected neurospheres (30 FFU/sphere) and glioblastoma cell lines (24 hpi). The latter was performed under low (experiment 1) and high (experiment 2) cell density conditions). Differentially secreted proteins were defined by pairwise t-test (p ≤ 0.05) or ≥2-fold changes in all replicates (neurospheres only). Differentially secreted proteins derived from experiments with SNB-19 and U87-MG cells, respectively, were grouped for further analyses. **(A-C)** Volcano plots showing secretome changes of neurospheres (A, n = 2, 128 proteins), U87-MG cells (B, experiment 1, n = 4, 259 proteins) and SNB-19 cells (C, experiment 2, n = 2, 192 proteins). Differentially secreted proteins are highlighted in blue (reduced) or red (increased). Altered proteins relevant for neural cell migration or neurite outgrowth are labelled and highlighted by black circles. Proteins affected in neurospheres and U87-MG cells are labelled in bold letters. Viral proteins (S = structural,NS = non-structural) are labelled. **(D)** Comparison of biological processes enriched among differentially secreted proteins (GOBP, Fisher exact test, p ≤ 0.05, 10 most enriched processes per model). Results were clustered for visualization (slim semantic similarity settings). Significance levels are reflected by the size of bubbles. Processes potentially related to neural cell migration or neurite outgrowth are highlighted in blue. **(E)** Proteins differentially secreted by neurospheres or both glioblastoma cells (viral proteins excluded). Fold changes are reflected by colours (blue = decreased, red = increased abundance in the secretome). Proteins differentially secreted by neurospheres or glioblastoma cells exclusively are depicted in black or grey letters, respectively. Proteins differentially secreted by neurospheres and U87-MG cells are highlighted in blue. **(F)** Proteins relevant neural cell migration or neurite outgrowth with significantly altered abundance in the whole cell proteome (see Figure 4) or secretome across all experiments (70 proteins). Fold changes induced by infection are reflected by colours (blue = decreased, red = increased abundance). Proteins with significantly altered abundance in the proteome or secretome of neurospheres are highlighted in purple and green, respectively. Proteins affected in neurospheres and glioblastoma cells (any experiment) are labelled with stars.

In total, we identified 167 proteins differentially secreted upon infection, with balanced up- and down-regulation upon VEEV infection (Supplementary Fig. 5H-I). Of the 42 most differentially secreted proteins, only three were affected by serum-depletion on whole cell proteome level, indicating that the majority of the observed secretome changes were not caused by serum depletion, but indeed by VEEV infection (Supplementary Fig. 5J). Of note, 187 proteins were exclusively quantified in the secretome, but not in the intracellular proteome and 114 of all differentially secreted proteins showed no detectable changes on whole cell proteome level (Supplementary Fig. 5H).

When comparing secretome changes of neurospheres and cell lines after VEEV infection, we observed model-specific alterations, in line with the differential intracellular proteomes (Fig. 5D, Supplementary Fig. 5I). Of the 167 proteins affected across all experiments, only six proteins were affected in more than one model, none of them in all three models (Fig. 5E). However, we identified 2 proteins with significantly reduced secretion in both, neurospheres and U87-MG cells, namely the phospholipid transfer protein (PLTP), and amyloid-beta precursor protein (APP), a protein with neurotrophic function when secreted^30^.

In line with the previously observed reduced cytoskeletal dynamics on intracellular level, we found evidence for disturbed neuron projection in all three models (Fig. 5D, Supplementary Fig. 5K-M). VEEV infection of neurospheres reduced the level of secreted microtubule-associated protein 1B (MAP1B) 7-fold and that of secreted MDK 20-fold. Similar to APP, secreted MAP1B acts neurotrophic^31^. MDK is a cytokine and growth factor released mainly by glial cells that promotes neuronal migration and embryonic neuron survival^32, 33^.

The potential impact of VEEV infection on neurite outgrowth was further highlighted by identification of, in total and across all experiments, 70 proteins that are related to the process of neuron projection. These proteins showed altered abundance on proteome or secretome level upon infection with the majority being downmodulated (Fig. 5F). Considering proteins affected in more than one experiment, VEEV reduced expression and/or secretion of 65 of these proteins (compared to 24 proteins with increased abundance).

To further integrate secreted factors with intracellular signaling pathways, we manually performed upstream regulator analysis based on differentially expressed proteins in VEEV-infected neurospheres. We linked MDK and APP to candidate cell-surface receptors and downstream intracellular effectors involved in cytoskeleton remodeling and neurite outgrowth (Supplementary Fig. 6A, Table S9-10). Notably, network reconstruction revealed convergence of these signaling routes on key regulators of actin and microtubule organization, consistent with the observed impairment of neurite outgrowth. In line with this, an unbiased upstream regulator analysis restricted to cellular proteome changes identified APP as one of the most significant upstream regulators, with 23 targets detected among the differentially expressed, intracellular proteins (Supplementary Fig. 6B–C). Together, these results suggest that VEEV-induced alterations in the secretome are functionally connected to intracellular pathways controlling cytoskeletal dynamics and neuronal projection.

In conclusion, secretome profiling uncovered infection-induced downmodulation of cytokines and growth factors. Our data point toward a VEEV-induced impairment of paracrine effects of glial cells including reduced MDK, APP, and MAP1B secretion that contributes to reduced neurite outgrowth.

## Discussion

Here we combined human iPSC-derived neurosphere technology with multi-omics and neurite outgrowth analyses to uncover whether and how VEEV infection impairs neuronal function. We found that human neurospheres, which express markers of neural progenitor cells, neurons and glial cells, are permissive to VEEV. Strikingly, infection impairs neurite outgrowth. To unravel the underlying mechanism, we demonstrated that VEEV downmodulates cytoskeletal pathways including Rho GTPases by inducing a translational shutoff. Moreover, VEEV lowered release of the neurite outgrowth factors MDK, APP, and MAP1B.

The clinical observation that VEEV infection can impair brain development, prompted us to investigate possible underlying mechanisms using human iPSC-derived neurospheres. These complex 3D spheres have emerged as a powerful, reductionist model for investigating human brain development *in vitro*. In our study, we confirmed the heterogeneous neural cell composition by transcriptomic, proteomic, and immunocytochemical analyses. Using an adapted version of the 2D-NIM protocol with differentiation in CINDA+ medium, as described by Hartmann et al. (2023)^13^, the neurospheres showed efficient neural commitment, expressing markers of neural progenitors as well as neuronal and glial lineages. Immunostaining for TUJ1 and GFAP confirmed the neuronal and astroglia populations, including astrocytes, radial glia and neural progenitor cells^34, 35, 36^. In line with previous forebrain-like organoid and neurosphere models^37^, our analyses support the presence of neural progenitor cells (SOX2, NESTIN, PAX6), immature neurons (DCX, TUJ1) and more mature neuronal and glial populations (MAP2, GAD1, GRIN1, TH, GFAP, S100B). In contrast to regionally patterned brain organoids with distinct ventricular-like zones and cortical organization^38, 39^, the neurospheres show an unstructured mixture of neural progenitor, neuronal and glial markers. This is consistent with early, pre-organized stages of CNS development and with reports on human neurospheres lacking clear cytoarchitectonic regionalization^18^. Together, these features are suggestive of a developing, forebrain-like identity rather than a more mature brain tissue and suggest that the neurospheres capture selected molecular key aspects of early brain development in vitro^38^. In addition, we observed migratory behavior in plated neurospheres, further supporting their use to study selected aspects of how neurotropic viruses impact the developing brain.

Leveraging this platform, we observed broad cell tropism of VEEV in human neurospheres targeting neurons and glial cells. This closely resembles infection patterns in small animal models. In mice, VEEV-TC83 infected neurons and glial cells after neuroinvasion independent of the peripheral injection route and regardless of the presence or absence of other cell types like microglia or oligodendrocytes^40, 41^. Interestingly, we observed infection kinetics of VEEV in neurospheres that were delayed in comparison to cell culture kinetics. In neurospheres titers increased until 48 hpi, while in cell culture titers typically plateau after 16 to 24 h. This suggests that VEEV either less efficiently infects neurospheres or that antiviral responses in neurospheres delay replication. We observed high expression levels of known VEEV host factors including the receptor LDLRAD3 and the replication factor FXR1, thus limited permissiveness is unlikely. In contrast, the ISGs IFITM3 and ISG15, which have proven anti-alphavirus activity^42, 43^, were upregulated on protein level in the neurospheres. Interestingly, the antiviral activity of ISG15 is visible in alphavirus infected murine brains, but not in isolated neuron cultures, further strengthening the notion that complex cellular networks are critical to understand virus-host interaction in the CNS.

While non-human primate- and rodent models described infiltration of immune cells and neuronal cell death, there is currently no intravital imaging study available, that would allow the assessment of neurite outgrowth upon VEEV infection. In contrast, we here describe direct visualization of neurite outgrowth dynamics upon VEEV infection in a controlled, human cell-based model. Considering the reports on memory loss and cognitive developmental impairment in VEEV patients, our human neurosphere model bridges a critical gap in mechanistically understanding how VEEV alters human CNS cell function in the developing brain. In addition, the model established here complements the recently developed co-culture spheroid system composed of iPSC-derived neurons and astrocytes^43^. The latter mimics more mature neuronal stages while lacking neural progenitor and radial glia populations, thereby modeling distinct aspects of adult brain infection and antiviral response.

Alphaviruses are well known to induce host translational shutoff to suppress antiviral protein synthesis and favor viral RNA translation. New World alphaviruses including VEEV shut down translation using their nsP2 and/or capsid proteins^44, 45^. In the neurosphere model, we confirmed translational shutoff upon VEEV infection. In particular, ISGs such as RIG-I were induced on mRNA, but not on protein level. Interestingly, we also observed a modest downregulation of protein expression of the VEEV host factor FXR1 across all three experimental models. Notably, FXR1 is not only a VEEV host factor, but also important for neurodevelopment^46^. The protein is strongly expressed in the developing brain and genome-wide association studies have linked it to neurological disorders. FXR1 knockdown leads to reduced neurite outgrowth^47^. Hence the here observed reduction of FXR1 likely not only keeps VEEV replication in check but also contributes to the impairment of neurite outgrowth.

Only few studies have addressed the impact of neuropathogenic viruses on neurite outgrowth. Herpesviruses, which cause persistent infection, upregulate neurite outgrowth by reducing expression of repellents of neurite outgrowth^48^ and by upregulating secretion of neurotrophic proteins^49^. In contrast, RNA viruses such as borna disease virus (BDV) and tick-borne encephalitis virus (TBEV) inhibit neurite outgrowth by preventing release of the outgrowth factor amphoterin^50^ and by impairing Rho GTPase pathways, respectively^51^. Interestingly, our multi-omics approach uncovered that VEEV, which is a zoonotic RNA virus like BDV and TBEV, impaired both, the secretion of outgrowth factors and RhoGTPase pathways. Consistent with this, a recent in vivo study demonstrates that VEEV infection also perturbs Rho GTPase–associated and synaptic signaling pathways in the hippocampus, linking these molecular alterations to long-term neuronal dysfunction and behavioral deficits^11^.Further, human neurosphere data support the view that Ras homolog family member A (RhoA)-related signaling acts as a functionally broad neurodevelopmental hub, with perturbation affecting multiple key processes, including neurite outgrowth and radial glia migration^52^. Rho kinases and their downstream effectors are essential for actin remodeling and thus neurite extension, axon guidance, and the migration of neurons along radial glia. Specifically, Rho GTPases phosphorylate targets like LIMK and cofilin to regulate growth cone dynamics^53, 54^. Pathway enrichment analyses of proteomic data revealed that VEEV reduces levels of cytoskeletal regulators, including DST and multiple Rho GTPase effectors. This indicates a VEEV-induced disruption of neuronal morphogenesis and migration, two processes critical for brain development^55, 56, 57^. In animal models, impairment of these pathways destabilizes growth cones, restricts neurite outgrowth, and disrupts cortical layering, ultimately leading to cortical malformations and defective neural circuits with lasting consequences for brain function and connectivity^58, 59^. This supports the interpretation that VEEV-induced suppression of Rho GTPase-associated pathways may have broader consequences for neuronal morphogenesis than neurite outgrowth defects alone. In contrast to the above mentioned BDV and TBEV studies, we here used a complex 3D culture of neurons and glial cells, which allowed us to not only assess neuron-intrinsic mechanisms, but also effects of neighboring glial cells.

Secretome analyses allow to elucidate intercellular communication that often remains concealed by conventional -omics approaches. At the same time, secretome profiling by LC-MS remains technically challenging due to the generally low abundance of secreted proteins and the potential interference of medium components. When applied to infection experiments with cytopathic viruses such as VEEV, secretome analyses are complicated by the limited time frame for accumulation of secreted proteins. Here we controlled for these confounding factors and revealed that VEEV infection of neurospheres downmodulates secretion of at least three proteins, namely MDK, APP, and MAP1B, which directly impact neurite outgrowth. MDK is expressed in the developing brain and stimulates neurite outgrowth^60^. Midkine is actively investigated as a promising neurotrophic therapy for perinatal brain injury^61^, suggesting its downregulation as a contributor to VEEV-mediated neurite defects. APP in its soluble form facilitates neurite outgrowth by inhibiting Wnt signaling^62^. Here, we observed VEEV-induced modulation of Wnt signaling pathways and reduced APP secretion, suggesting this contributes to the observed neurite outgrowth deficiency. Given the established neurotrophic and neuritogenic functions of soluble APP, its downregulation upon VEEV infection may not only contribute to pathogenesis but also highlight APP signaling as a potential target to restore VEEV-associated neurite outgrowth inhibition^63, 64^. In line with this, we observed neurite outgrowth defects not only in VEEV infected, but also in bystander neurons, indicating that reduced secretion of neurite outgrowth factors from VEEV infected cells also has paracrine effects. Finally, we observed reduced extracellular levels of MAP1B, which is an intracellular microtubule-associated protein that stimulates neurite outgrowth^65^. MAP1B is released from neurons within exosomes^66^, hence we detected MAP1B in the secretome fraction. Altogether, these findings underline the VEEV double whammy on neurite outgrowth in neurospheres, downmodulating not only cytoskeletal pathways, but also cytokine release.

Even though neurospheres effectively address key features of human brain development, some limitations must be considered. Our neurospheres lack oligodendrocytes, microglia, and additional immune cell populations, excluding their specific contributions to pathogenesis such as myelination defects, synaptic pruning, and inflammatory responses. While we could demonstrate robust neuron-astrocyte-driven innate immune activation, the absence of microglia precludes modeling of their specialized role in virus-host interactions. Additionally, fatty acid metabolism dysregulation noted in our omics data suggests metabolic consequences that may require the integration of oligodendrocytes to fully address influences of VEEV infection on myelin formation^67, 68^. Another limitation stems from the inability to define the precise cellular origin of secreted factors in bulk secretome preparations — single-cell secretomics approaches are thus a vital direction for future work.

Collectively, our data demonstrate that human iPSC-derived neurospheres are a relevant and informative platform to study VEEV neuropathogenesis and the impact of infection on developing neural tissue. While reductionist in nature, the model captures key features of the disease process and provides mechanistic insight into virus-induced disruption of neurite outgrowth and host cytoskeletal signaling. Importantly, this platform can be further expanded to address more complex developmental and functional endpoints, including neuronal network formation^69, 70^, thereby deepening mechanistic understanding and supporting future antiviral strategy development.

## Methods

### Maintenance of Baby Hamster Kidney (BHK) and glioblastoma cells

All cells were maintained in a humidified incubator at 37 °C and 5% CO_2_. Baby Hamster Kidney^71^ (BHK, kindly provided by C. Goffinet) and glioblastoma cell lines^72, 73^ (kindly provided by P. Claus (SNB-19) and O. Keppler (U87-MG)) were cultured in 10 cm dishes and subcultivated every 3-4 days at confluency of approximately 80%. BHK cells were cultured in DMEM (Gibco) with 10% fetal calf serum (FCS, Capricorn scientific), 0.1 mM non-essential amino acids (Gibco) and 2 mM L-Glutamine (Gibco). The glioblastoma cell line SNB-19 was cultured in high-glucose DMEM with 10% FCS and 1% Penicillin/ Streptomycin (Thermo Fisher, 10.000 U/ml). The glioblastoma cell line U87-MG was cultured in high-glucose DMEM (Thermo Fisher) with 10% FCS, 1% Penicillin/ Streptomycin, 1% L-glutamine and 1% sodium pyruvate (Thermo Fisher). Mycoplasma testing was performed twice a year applying the Mycoplasma Detection Kit (Southern Biotech).

### Maintenance of induced pluripotent stem cells (iPSCs)

The human induced pluripotent stem cell (iPSC) line IMR90-04, originally obtained from WiCell^74^, was used for the generation of neural progenitor cells and neurospheres. Monthly Mycoplasma testing was performed, applying the Mycoplasma PCR Detection kit (Applied Biological Materials). IMR90-04 cells were cultured in StemMACS iPS-Brew XF medium (Miltenyi) on Matrigel (growth factor reduced; Corning) -coated 6-well plates. Cells were passaged with 0.02% ethylenediaminetetraacetic acid (EDTA) in phosphate buffered saline (PBS) without Mg^2+^ and Ca^2+^ when they reached confluence of 70-80%. After thawing and subcultivating of the iPSCs, 10 μM Rho-Kinase inhibitor Y-27632 (Target Mol) was supplemented to the culture medium until the next medium change.

### Neurosphere formation

Induction of neural progenitor cells derived from IMR90-04 cells and the subsequent formation of neurospheres was adapted from Fritsche and coworkers^13^. Briefly, 2 × 10^6^ IMR90-04 cells, dissociated using Accutase (Sigma-Aldrich), were seeded per well in a 6-well plate previously coated with 0.1% polyethyleneimine (PEI) (Sigma-Aldrich) and 15 µg/mL laminin (Sigma-Aldrich). Cells were cultured in 2D-neural induction medium (NIM) (Table S1) supplemented with 10 μM SB-431542 hydrate (MedChemExpress/Hycultec) and 0.5 μM LDN-193189 hydrochloride (Sigma Aldrich) for the first 11 days, with daily medium changes. From day 12 to 21, NIM was supplemented with 20 ng/mL of basic fibroblast growth factor (bFGF; R&D) only, also with daily medium changes. On days 12 and 17, cells were detached using Accutase and reseeded at a density of 2 × 10^6^ cells per well in NIM containing 20 ng/mL bFGF and 10 μM Y-27632. Y-27632 was removed with the subsequent medium change. On day 21, neural progenitor cells were dissociated using Accutase and cryopreserved in NIM supplemented with 10% DMSO, 20 ng/mL bFGF and 10 μM Y-27632.

For 3D neurosphere formation, approximately 1.3 × 10 neural progenitor cells were thawed and seeded per well in NIM supplemented with 20 ng/mL bFGF and 10 μM Y-27632, on PEI- and laminin-coated 6-well plates. After four days of daily medium exchange with NIM containing 20 ng/mL bFGF and 10 μM Y-27632, cells were detached using Accutase and resuspended in NIM supplemented with 20 ng/mL bFGF and 10 μM Y-27632. Approximately 1 × 10 cells in 3 mL were transferred per well into anti-adherence solution-treated wells (Stemcell Technologies) of a 6-well plate. The plates were placed on an orbital shaker at 140 rpm, 19 mm diameter (MaxQ 2000 CO Plus, Thermo Fisher Scientific) in the incubator to promote neurosphere formation from single cells over seven days. On day 8 of neurosphere formation, the spheres were chopped using a McIlwain mechanical tissue chopper to a size of 0.25 mm. The sphere fragments were transferred to CINDA+ medium (Table S2) freshly supplemented with 300 μM dibutyryl cyclic AMP sodium salt (Sigma Aldrich), and orbital shaking was continued. Spheres were differentiated for two weeks in CINDA+ medium before another chopping step was performed as described above. Throughout the differentiation process, a 50% medium change was performed twice per week. After a total differentiation time of four weeks, the spheres were used for experiments.

For experiments with adherent spheres, single spheres were placed on 50 µg/mL poly-L-ornithine (PLO) (Merck) and 15 µg/mL laminin-treated 96-well plates (Greiner-cellstar clear F-bottom) 24 h before experiment was performed and were cultivated in 100 µL CINDA+ medium.

### Gene expression analysis during differentiation of iPSCs to neurospheres by qPCR

Cells were cultured and harvested as in the main manuscript. RNA isolation, reverse transcription and SYBR green-based gene expression analysis was performed as described in Schenke et al.^75^. RT-qPCR was performed on a Biorad cfx96 touch cycler. Gene expression levels were calculated by the 2^-ΔΔC^ method^76^. Gene expression of target genes was normalized on expression of three stably expressed reference genes cyclophilin A (*PPIA),* 60S acidic ribosomal protein P0 *(RPLP0),* ribosomal protein S23 (*RPS23)*. The respective primer pairs for detection of the genes of interest are shown in Table S8. The genes analyzed were *OCT4* (pluripotency), *SOX2* (neural stem cell identity), *PAX6* and *NESTIN* (early neural specification and neural progenitors), *DCX* and *TUJ1* (immature neurons and early neuronal differentiation), *MAP2* (neuronal maturation), *GAD1* and *GRIN1* (inhibitory and excitatory neuronal phenotypes), *TH* (dopaminergic lineage specification), and *GFAP* and *S100B* (astroglial markers).

### Generation of virus stocks

The VEEV TC-83 infectious clone expressing GFP under a subgenomic promoter was kindly provided by Ilya Frolov^77, 78^. 15 µg of plasmid DNA was linearized using MluI (NEB) and extracted with the QIAprep Spin Miniprep Kit (Qiagen) according to the manufacturer’s instructions. For *in vitro* transcription, 2 μg of purified DNA were pre-incubated in a 100 µL reaction mix with final concentrations of 1× RNA polymerase buffer (NEB), 25 mM per rNTPs (Roche), 5 mM Ribo m^7^G Cap Analog, 40 U/µL RNasin® Ribonuclease Inhibitor and 20 U/µL SP6 RNA polymerase (all Promega) for 2 h at 37 °C. Transcription was performed by adding 4 µL of SP6 RNA polymerase (Promega, 20 U/mL) for 2 h at 37 °C. Viral RNA was purified using the NucleoSpin RNA Clean-up kit (Macherey-Nagel) according to the manufacturer’s recommendations. 5 × 106 BHK-21 cells were transfected with 10 µg of viral RNA in 500 µL OptiMEM using a Gene Pulser Xcell modular electroporation system (Bio-Rad). 1 × 10^7^ transfected cells were pooled and cultured in cell culture flasks in 10 mL cell culture medium for 48 hpi. Supernatant was harvested followed by centrifugation at 195 × g for 5 min and filtered through a 0.45 μm pore size filter. Aliquots of virus stocks were stored at -80 °C.

### Quantification of infectious viral particles by titration

TCID_50_ endpoint dilution assays were used to measure viral titers. Cells were seeded in 96-well plates at a density of 1 × 10 SNB-19 (virus stock SNB-19 infections, neurospheres’ supernatants) cells or 2 × 10 U87-MG (virus stock U87-MG infections) cells per well in 90 µL medium and incubated at 37 °C with 5% CO for 24 h. Virus samples were pre-diluted 1:10 in the respective medium. 10 µL were added to the first well followed by careful mixing, followed by a series of ten consecutive 1:10 dilutions using fresh pipette tips for each transfer. Each sample was assayed in 4–6 replicates, with at least one well per plate serving as a negative control. GFP-positive wells were quantified 24 hpi by fluorescence microscopy. TCID_50_/mL were calculated by the Spearman–Kärber method using an Excel-based tool provided by Marco Binder (V1; updated version available at Heidelberg University Medical Center, Department of Infectious Diseases).

### Quantification of viral genome copy numbers by qPCR

Cells were cultured, infected and harvested as described above. Total RNA was extracted from cell pellets using the Nucleospin RNA Kit by Macherey-Nagel according to the manufacturer’s instructions (see Table S5 for RNA yields). RT-qPCR was performed on a Roche LightCycler using a TaqMan assay (Applied Biosystems AgPath-ID One-Step RT-PCR Reagents) targeting the viral genome and the reference gene ACTB in a 96-well plate format (for details including primer sequences see Tables S6 and S7). For each reaction, 2.5 µL of isolated RNA (corresponding to 6.25% of total isolated RNA) was used. Standard curves were generated from samples with defined copy numbers of synthetic viral RNA (VEEV 5’UTR to polyA tail), which were derived from a serial 1:10 dilution and ranged from 5 × 10^1^ to 5 × 10^8^ copies per microliter. All reactions were performed in technical triplicates and included no-RT controls. Viral RNA copy numbers in neurosphere samples were calculated based on standard curves and normalized to the total RNA content per sample.

### VEEV infection

For infection of floating neurospheres, spheres were cultured on 6-well plates with 30 spheres and 2 mL CINDA+ medium per well. To achieve infectious doses of 3, 30 or 300 FFU per sphere, a respective amount of the virus stock was added to every well to be infected, ending up with final dilutions of the virus stock between 10^-3^ and 10^-5^. Of note, a calculation of MOIs was not possible due to the varying number of cells per sphere, however, we expected approximately 1,500 cells per sphere which corresponds to MOIs of 0.002 – 0.2. Spheres were washed twice with PBS 4 hpi followed by adding fresh culture medium. For infection of plated spheres, a single sphere per 96-well in 100 µL CINDA+ medium was infected following the same procedure except for the applied infectious doses (5, 45 or 450 FFU per sphere) and leaving out the washing step 4 hpi. For infection of glioblastoma cells, 1 × 10^5^ (SNB-19) or 3 × 10^5^ cells (U87-MG) were seeded on 12-well plates and infected 24 h post plating with an MOI of 0.1 (whole cell proteome and secretome). Cells were washed twice with PBS 4 hpi followed by adding fresh culture medium. For upscaled secretome experiments, 0.7 × 10^6^ (SNB-19) or 1.4 × 10^6^ cells (U87-MG) were seeded on 6-well plates followed by the same infection procedure. For all experiments, successful infection was verified by GFP expression via fluorescence microscopy before harvesting cells between 24 and 72 hpi (Supplementary Figure 2C). Uninfected samples served as controls and were treated the same way.

### Fluorescence Imaging (overview images)

Fluorescence images of VEEV-infected neurospheres were acquired using an EVOS™ M5000 Imaging System (Thermo Fisher Scientific). GFP signal was detected using the EVOS GFP light cube (excitation 470 nm, emission 525 nm). Images were captured using identical exposure settings within each experiment to allow qualitative comparison of infection intensity across samples. Brightfield and fluorescence images were acquired sequentially at 4× magnification.

### Fixation of neurospheres

4% paraformaldehyde (PFA) in PBS was added to the culture medium to achieve a final PFA concentration of 2%. After 10 min incubation at RT, complete medium was removed and 4% PFA in PBS was added followed by a second incubation for 10 min at RT. Cells were washed twice with PBS and stored in PBS at 4 °C after sealing with parafilm. The procedure was carried out with great caution to leave cell morphology intact.

### Cell viability (CTB) and cytotoxicity (LDH) assays

Single neurospheres were placed on PLO/laminin-treated 96-well plates as described above. For lysis controls, 0.2% Triton X-100 was applied 45 min before start of the respective assay. Untreated spheres in CINDA+ medium served as controls. Cytotoxicity was assessed by quantifying lactate dehydrogenase (LDH) release into the culture medium using the CytoTox-ONE™ Homogeneous Membrane Integrity Assay (Promega,). 50 µL of respective medium were transferred to an empty well of a 96-well plate (Sarstedt, 96 Flat Transparent). 50 µL of CytoTox-ONE™ reagent (RT) were added, followed by incubation for 2 h at room temperature. Cellular viability was quantified with the CellTiter Blue Cell Viability Assay (Promega). 50 µL of respective medium was incubated with 83 µL of CTB working solution (CTB working solution = CTB reagent 1:7.5 diluted with CINDA+), followed by incubation for 2 h at 37 °C. Read outs were performed with a Spark microplate reader (TECAN, excitation: 550 nm, emission: 620 nm). Experiments were performed with 3-5 technical replicates per plate. Biological replicates were derived from experiments performed at different days and with spheres derived from independent differentiations.

### Neurite outgrowth quantification

Spheres were placed on PLO/laminin-treated 96-well glass plates (Ibidi) as described above. The spheres were cultured for 24 h to allow attachment to the bottom of the plate and initial neurite outgrowth. The length of outgrown neurites was documented before treatment of spheres (24 h post plating) by bright field microscopy (Nikon, Eclipse Ti). Subsequently, cells were infected with VEEV as described above using uninfected cells as controls. Cells treated with 10 µM PP2 (Merck), a known inhibitor for neurite outgrowth/neural progenitor cell migration, served as a positive control for inhibition of neurite outgrowth. DMSO treatment (0.1%) was used as a solvent control for PP2 treatment. 24 h post treatment, spheres were fixed with 4% PFA and imaged a second time. The length of the same selected neurite before and after treatment was measured in ImageJ with the straight-line tool. Each neurite was traced from the edge of the sphere core in a straight line to its distal end. To determine the effect of the treatment on neurite outgrowth, the neurite lengths measured after treatment were normalized to neurite lengths measured before treatment (ratio). A total of five individual experiments were conducted with spheres gained from five independent differentiations initiated by two individual iPSC and resulting neural progenitor batches. Up to seven spheres per experiment and four neurites per sphere were analyzed. Spheres with abnormal morphology, indicating reduced overall cell viability, occurred across all experiments and were not taken into account for the experimental read out.

### Confocal Microscopy

Both, floating and plated spheres, were infected, fixed and prepared for immunofluorescence imaging in 96-well plate format using a single sphere per well. Primary antibodies (Table S3) were diluted in 10% goat serum (Sigma Aldrich) in PBS with 0.05% Tween 20 (PBST) and incubated overnight at 4 °C. After three washing steps with PBS, cells were incubated with fluorophore conjugated secondary antibodies diluted as indicated in Table S3 in 10% goat serum in PBST for 60 minutes at RT in the dark. The secondary antibody was removed, and the samples were washed with PBS three times. The floating spheres were collected and transferred onto a microscopy slide. After mounting (ProLong Gold antifade reagent with DAPI, Invitrogen), the spheres were covered with a cover slip and stored in the dark until imaging. The adherent spheres were incubated with Hoechst (5 µg/mL) for 10 minutes at RT to stain DNA. After washing with PBS, the spheres were stored in PBS until imaging.

Fluorescence imaging was performed using a LSM Meta 800 confocal laser scanning microscope (Zeiss). Excitation was achieved with diode lasers at 493 nm for GFP and Alexa Fluor 488, 577 nm for Alexa Fluor 568, 653 nm for Alexa Fluor 647, and 353 nm for DAPI or Höchst, respectively. Emission was collected using the standard spectral detectors of the system with appropriate bandwidth settings for each fluorophore. Images were acquired with exposure settings adjusted to maximize signal while avoiding pixel saturation Representative replicates were selected for image processing. Raw microscopy images were acquired as 16-bit.czi files. For figure preparation, individual fluorescence channels were first converted to RGB using Fiji^79^. After RGB conversion, brightness and contrast were manually adjusted using the “Brightness/Contrast” tool for each channel to optimize signal visibility. Channel-specific display ranges (min/max) were set to reduce oversaturation in highly fluorescent channels (e.g., green, GFP) and to ensure that weaker signals (e.g., red, AF 568) remained clearly visible in the merged images. All adjustments were linear and applied exclusively for visualization purposes. No gamma correction or nonlinear enhancements were used. Display settings for each image and channel are listed in Supplementary Table S4.

### Sample preparation for transcriptome, whole cell proteome and secretome analyses

Cells were cultured and infected with VEEV as described above. Biological replicates were derived from experiments performed at different days and, in case of neurospheres, with spheres derived from independent differentiations. For transcriptome and whole cell proteome analysis under standard conditions, medium was replaced by the respective maintenance medium after the washing step during the infection procedure. For secretome analysis as well as transcriptome and whole cell proteome analyses under serum-free conditions, maintenance medium was replaced by medium lacking B27+ (neurospheres) or FCS (SNB-19 and U87-MG), respectively. Stem cells were cultured in TeSRE8 (Stemcell Technologies) for 3 days for both, whole cell proteome and secretome analysis. Conditioned medium was transferred into 2 mL reaction tubes 24 hpi (neurospheres, glioblastoma cells) or 3 days post plating (iPSCs) for subsequent secretome analysis. The medium was spun down at 2000 × g and 4 °C for 5 min to get rid of cell debris. Supernatant was transferred again to a fresh 2 mL reaction tube and frozen at -80 °C till further processing. Remaining cells were washed twice with PBS. Adherent cells were detached by adding trypsin for 5 min at 37 °C and transferred into a 1.5 mL reaction tube. Cells were spun down at 300 × g and 4 °C for 5 min, washed with PBS once and spun down a second time. Supernatant was discarded and cells were stored at -80 °C till further processing. Cells cultured in suspension were transferred into 1.5 mL reaction tubes during the second PBS washing step, followed by centrifugation, removal of the supernatant and storage at -80 °C. While different samples were used for the transcriptomics and proteomics experiments, samples from the same serum-free cultures were employed for both the proteome and secretome studies.

### Transcriptome analysis by RNA sequencing

RNA for transcriptome analysis was isolated from cell pellets using the NucleoSpin RNA Clean-up kit (Macherey-Nagel) according to manufacturer’s instructions. Library preparation, RNA sequencing as well as raw data processing and initial data analysis was performed by Novogene (Beijing, China). In brief, mRNA was purified from total RNA with poly-T oligo-attached magnetic beads and fragmented followed by cDNA synthesis with random hexamer primers. Construction of the sequencing libraries comprised end repair, A-tailing, adapter ligation, size selection, amplification, and purification. The library quality control entailed quantification and size distribution analysis. Libraries were pooled and sequenced on an Illumina platform. Raw sequencing reads were processed with in-house scripts to remove adaptor sequences, poly-N regions, and low-quality reads. Clean reads were aligned to the Homo sapiens reference genome (Ensembl GRCh38.p12, accession GCA_000001405.27) using HISAT2 v2.0.5. In addition, the sequence corresponding to NCBI accession L01443.1 was downloaded from the NCBI nucleotide database for a separate alignment. Transcript assembly was performed with StringTie v1.3.3b. Gene expression levels were quantified using featureCounts v1.5.0-p3 to obtain read counts per gene, which were then normalized to FPKM (fragments per kilobase of transcript per Million mapped reads) based on the length of the gene and reads count mapped to this gene. A principal component analysis (PCA) was used to assess data quality and biological variability. Samples with poor data quality were excluded from further analyses. Differential expression analysis was performed using the DESeq2 R package (v1.20.0) and Benjamini–Hochberg correction for p value adjustment. Enrichment analysis was done with the clusterProfiler R package to determine enrichment among differentially expressed genes with adjusted p values ≤ 0.05 taking into account Gene Ontology (GO) terms and Kyoto Encyclopedia of Genes and Genomes (KEGG) pathways.

### Whole cell proteome and secretome analyses by liquid chromatography-mass spectrometry

For whole cell proteome analysis, proteins were purified from cell pellets and processed as described recently^80^. In brief, cells were lysed with RIPA buffer followed by acetone precipitation of proteins. Protein amounts were determined by BCA assay and 20 µg of protein per sample were digested in solution using dithiothreitol (DTT) for reduction of disulfide bridges, iodoacetamide (IAA) for alkylation of thiols and trypsin gold (Promega) for cutting proteins into peptides. For secretome analyses, 400 µL of conditioned medium was mixed with 1600 µL ice-cold acetone followed by incubation at -20 °C for 15 h (overnight). Precipitated proteins were spun down at 12.000 g and 4 °C for 15 min and supernatant was discarded carefully. The protein pellet was dried at RT for 30 min, resuspended in 50 µL 2 M Urea and 50 mM Tris-HCl (pH 8) and sonicated for 3 min. An aliquot of 10 µL was taken to determine protein concentration by BCA assays. The procedure was repeated with a higher amount of purified protein, if the protein amount was below detection limit. An aliquot of 20 µg protein in 50 µL 2 M Urea in 50 mM Tris-HCl (pH 8) was used for the tryptic digest.

0.5 µL DTT were added to a final concentration of 10 mM followed by 1 h at 37 °C. 5.5 µL IAA were added to a final concentration of 15 mM followed by 30 min at RT and protected from light. 5.6 µL thiourea were added to a final concentration of 100 mM. 500 ng (0.5 µL) trypsin were added followed by 15 h incubation at 37 °C. At the end of the tryptic digest, samples were acidified by addition of 50 µL 5% acetic acid to stop the enzymatic digest and to allow subsequent sample clean up. Digested peptides from both, whole cell proteome and secretome samples, were purified using self-made StageTips^81^ followed by resuspension in 40 µL of MS sample buffer (0.1% formic acid and 5% acetonitrile in MS grade water) and sonication for 3 min.

LC-MS runs were performed with a Vanquish Neo nanoflow UHPLC System coupled to a Nano Spray Flex Ion Source with stainless steel emitters (40 mm, OD 1/32) and an Orbitrap Eclipse mass spectrometer (Thermo Scientific) as described previously^80^. In brief, 500 ng peptides per sample were separated on a reversed-phase nanoViper™ PepMap™ column (150 mm length, Thermo Scientific™) applying an 80 min gradient with buffer A (5% ACN, 0.1% FA) and B (80% ACN, 0.1% FA) at 40 °C and a flow rate of 250 nL/min. Precursor scans were performed in the Orbitrap mass analyzer. 20 most intense precursors were selected for HCD fragmentation in the linear ion trap. Fragment ion scans were performed in the best performing analyzer at the time of the experiment, which was the Orbitrap analyzer for the whole cell proteome of VEEV-infected and uninfected neurospheres (24 hpi) and the ion trap analyzer for all other samples. MS2 scans in both analyzers were performed with normal mass range, maximum injection time set to auto, AGC target set to standard and data type set to centroid. MS2 scans in the Orbitrap were performed a resolution of 15,000, scans in the Ion trap were performed with a scan rate set to rapid.

Processing of LC-MS derived raw data was performed using MaxQuant software (V. 2.4.14.0)^82^. The implemented Andromeda search engine was applied for the identification and quantification of proteins. MS spectra were searched against UniProt database of human proteins (ID UP000005640, reviewed, database downloaded on 04/03/2024, taxonomy ID set to 9606) and VEEV proteins (strain Trinidad donkey, ID UP000146452, downloaded on 04/03/2024, taxonomy ID set to 11038) applying preconfigured settings (PSM and protein FDR = 0.01, min peptides = 1, min. unique peptides = 0, Match between runs disabled). In addition, Label-free quantification (min ratio count = 2), iBAQ quantification and writing the mzTab table was enabled.

### Omics data analyses

RNAseq data were analyzed by Novogene as described above and resulted in matrices containing the number of reads mapped to the respective reference genome, identified transcripts and the respective number of reads, normalized mean fpkm intensities, fold changes and (adjusted) p values derived from pairwise comparisons as well as biological processes enriched among differentially expressed genes. Further data analysis was restricted to filtering out results of interest. Gene expression profiles (like expression of neuronal markers) were restricted to transcripts with fpkm values ≥ 1. Applied thresholds for (adjusted) p values are described in the results section and figure legends.

Processed MS raw data were further analyzed based on the proteinGroups.txt file using Perseus Software (V. 1.6.15.0) ^83^ with standard setting unless otherwise specified. The term protein is used here instead of protein group. In general, data analysis was performed separately for selected subgroups of the whole dataset and identification of proteins affected by VEEV infection was based on pairwise comparisons of infected and uninfected samples and performed separately for each condition. iBAQ values were used to rank protein abundance within an experimental group, LFQ values were used to compare protein abundance across groups. iBAQ and LFQ intensities were log2-transformed. Additional transformation is stated in the figure legends, if applied. Data integrity was assessed by comparing the number of identified proteins, by the overall data distribution and by principal component analyses. Samples with poor data quality (significantly less identified proteins compared to other replicates of the same group) were excluded from further analysis. From the matrix of all identified proteins, common contaminants, proteins only identified by site modifications and proteins identified in the database of reverse protein sequences were removed (high-confidence filter). Furthermore, for analyses of neurosphere-derived samples, B27 medium components were removed, including catalase, (pro-) epidermal growth factor, insulin, neurotrophin-3, interferon-gamma, superoxide dismutases and (sero- / lacto-) transferrin. Unless otherwise stated, all comparative analyses were restricted to equal numbers of replicates in each group (single replicates removed if required) and to proteins with valid values in at least 65% of replicates in at least one of the respective experimental groups. Differential protein expression analysis was performed by pairwise t-tests or one way ANOVA, respectively, as indicated in the figure legends. Immunoglobulins were considered contaminants derived from cell culture media and excluded from the list of differentially expressed proteins. For enrichment analyses, the frequency of annotations among differentially expressed proteins was compared to the frequency within the group of all detected proteins. 1D and 2D enrichment analyses was performed as described by Cox and Mann^84^. Annotations were added based on Majority Protein IDs using Perseus Annotation file downloaded on 03/25/2025. Heat maps were generated without preprocessing. VEEV proteins were removed prior to statistical and enrichment analyses.

### Bioinformatic processing of results derived from omics data analyses

Matching of RNAseq data with proteome data was achieved by matching of gene names. Redundancy of GO terms resulting from enrichment analyses were reduced using the web-based tool REVIGO^85^ with medium or slim semantic similarity settings for grouping of pathways as described in the figure legends. Reactome IDs were matched with their respective entry names using Reactome’s Complete Pathway List. Redundancy of Reactome terms was reduced based on parent and child IDs derived from Reactome’s Pathways Hierarchy Relationship. Both lists were downloaded from the Reactome database (https://reactome.org/download-data, 03/31/2025). Pathway names were abbreviated to save space, unique assignment to the database entry was based on the IDs. Proteins involved in neural cell migration or neurite outgrowth were defined by GO terms comprising strings “migrat”, “project”, “exten” or “guid”, together with strings “neuron“, „axon“ or „dendrit”. Proteins were classified as extracellular using the GOCC slim database entries *extracellular region / space / matrix / organelle* and *external encapsulating structure*. Apoptosis markers were filtered based on KEGG and Uniprot Keyword annotations “Apoptosis”.

### Deconvolution of RNAseq data

Cell type proportions in bulk RNA-seq samples were estimated by supervised deconvolution using fpkm values, the Deconomix^86^ algorithm and a harmonized human neural organoid single-cell RNA-seq atlas of human neural organoids as reference^15^. Harmonized cell type annotations were used to define the reference cell types, and the provided log1p-normalized single-cell expression values were used directly because raw counts were not available. Cell types not compatible with the applied differentiation protocol were excluded from the reference, including mesoderm-derived populations (e.g., endothelial cells, mesenchymal cells, and microglia) as well as additional lineages unlikely to be present under these conditions (e.g., choroid plexus, neural crest, oligodendrocyte precursor, and perivascular stromal cells). The reference was restricted to highly variable genes and intersected with the bulk RNA-seq dataset, yielding 3,533 shared genes for model training and application. Deconvolution models were trained and evaluated using a 70:30 train-test split, and performance was assessed by Pearson and Spearman correlations between estimated and expected cell type proportions. The trained models were applied to protein-coding bulk RNA-seq data restricted to the same gene set. Differences in inferred cell type proportions between Mock- and VEEV-treated samples were assessed using linear models and Wilcoxon tests with Benjamini-Hochberg correction. For cell type-associated transcriptional analyses, normalized bulk expression values were modeled gene-wise as a function of inferred cell type proportion and its interaction with VEEV status.

Further, we performed unsupervised cell-type deconvolution via non-negative matrix factorization, implemented in the R-package NMF^87^- For this purpose, (1) fpkm-values were transformed via log1p transformation, (2) restricted to the 10,000 most abundant genes, (3) centered sample-wise, (4) shifted to yield a non-negative data matrix with the lowest value equaling zero, and (5) re-scaled gene-wise to a mean of 1. Then, a NMF with a rank of 4 was performed on the transformed gene-expression data. Derived cell-compositions were normalized to proportions. In line with the supervised analysis, bulk expression values were modeled gene-wise as a function of inferred cell type proportion and its interaction with VEEV status. Corresponding pathway enrichment analysis was performed using F-values (multiplied with the sign of the corresponding coefficient) using the rank-based test of the cameraPR function implemented in the R package limma^88^. Gene sets were retrieved from GO and KEGG databases using the R package EnrichmentBrowser ^89^. Gene sets representing cell types were retrieved from ^90^.

### Upstream regulator analyses

Differentially secreted proteins with signal peptides (web-based tool “SignalIP”) annotated with “signaling receptor binding” (GOMF GO:0005102) were used as ligand anchors. Candidate receptors of secreted proteins were identified using all identified transcripts with fpkm values ≥ 1. Known interactors (web-based tool OmniPath) were filtered for proteins annotated with “Cell membrane”, “Transmembrane” and “Receptor” (Uniprot Keyword database). Differentially expressed cytoskeleton and neurite-related proteins and other kinases were filtered applying annotations as listed in supplementary tables 9 and 10. Differentially expressed transcription factors were filtered based on the list from Human TF census^91^ (∼1600 TFs)). Protein–protein interaction networks were generated using STRING (experimental and curated evidence only) and visualized in Cytoscape, retaining proteins with at least one interaction and receptors with at least one interaction to secreted proteins.

### General statistics

Biological replicates were derived from experiments performed at different days. All pairwise comparisons (infected vs. uninfected; treated vs. untreated) were based on samples derived from the same experimental batches and on equal numbers of replicates. Listed numbers of identifications refer to transcripts or proteins detected in at least 1 replicate of the conditions included in the respective analysis or section described (based on raw counts or intensities). Numbers of quantified transcripts or proteins refer to those that remained after data processing as described above and shown in the graphs. Differentially expressed transcripts and proteins were defined by different the p value or FDR thresholds as indicated in the figure legends, depending on the data quality of groups being compared. Statistical tests on data apart from proteomics or transcriptomics data were performed using Graph Pad Prism software (V10), namely by one sample t-tests and one-way ANOVA with Dunnet post-hoc tests. The respective tests and applied significance thresholds are described in the results section and figure legends.

## Supporting information

File S01 - qPCR Neurospheres.xsx

File S02 - Dataset Overview.xlsx

File S03 - RNAseq + LC-MS (Unprocessed).xlsx

File S04 - Models Compared.xlsx

File S05 - VEEV (0-48 hpi).xlsx

File S06 - VEEV 24 hpi (Transcr).xlsx

File S07 - VEEV 24 hpi (WCP).xlsx

File S08 - VEEV 24 hpi (Secr).xlsx

File S09 - SerumDeplet_CellDens.xlsx

File S10 - Hits Combined Neur. Outgr. (MS).xlsx

File S12 - Neurite Outgrowth

## Data availability

RNAseq data (raw and processed data, mapped against human genome) will be deposited to the ArrayExpress repository with the dataset identifier E-MTAB-15795. LC-MS data (raw data and database search results) will be deposited to the ProteomeXchange Consortium via the <u>PRIDE</u>^64^ partner repository with the dataset identifier PXD069109. Source Data are provided with this paper and processed data as illustrated in the figures are available in supplementary files (supplementary files 01-12).

## Contributions

Cirksena K^#^, Haiber LM^#^, Kühne BA, Hupfeld J, Duven M, Fischer A, Schenke M, Richter F, Weiß S, Altenbuchinger M, Seifert N, Zacharias HU, Konzok S, Seeger B^‡^, Gerold G^‡^ BS and GG equally conceived and supervised the study. KC (infection, viability, transcriptomics, proteomics, secretomics, data analyses) and LMH (iPSC and neurosphere cell culture, neurite outgrowth and confocal microscopy) contributed equally to the experimental work. BAK, JH, MD, AF and MS contributed to several experiments and data analyses. SK supervised confocal microscopy. SW and FR contributed to Ingenuity Pathway Analysis. MA, NS and HUZ contributed to the deconvolution of RNAseq data, analysis and writing of the manuscript. BS and GG provided financial support. KC, LMH, BS and GG wrote the manuscript. BAK, JH, MD, AF, MS, FR, SW, MA, NS, HUZ, SK reviewed and revised the manuscript. All authors approved the final version of the manuscript. BS and GG contributed equally and have the right to list themselves last in bibliographic documents.

## Acknowledgements

The authors thank Alexander Krybus, Alexander Prokscha and Laura Rehburg for excellent technical assistance. The Gerold lab thanks Ilya Frolov for providing the VEEV reverse genetic system. We thank Peter Claus for providing SNB-19 cells and Oliver Keppler for U- 87 MG cells. The Seeger lab thanks Ellen Fritsche and her lab members (IUF Düsseldorf, now SCATH) for generously providing us with the opportunity to learn the method for neurosphere generation.

## Funding

Mass spectrometry work was funded by the Deutsche Forschungsgemeinschaft (DFG, German Research Foundation) – project number 497694394 granted to GG. GG further received support from the German Ministry of Research, Technology and Space and the Lower Saxony Ministry for Science and Culture (Professorinnen Programm III), the Knut och Alice Wallenbergs Stiftelse. Parts of the study were funded by the German Ministry of Research, Technology and Space – grant number 01KI2311B, granted to BS.

## Notes

### Competing Interest Statement

The authors have declared no competing interest.

