## Supporting Information for "Encephalitic alphavirus infection reduces neurite outgrowth and downregulates Rho GTPase pathways in human iPSC-derived neurospheres"

**Fig. S1:**
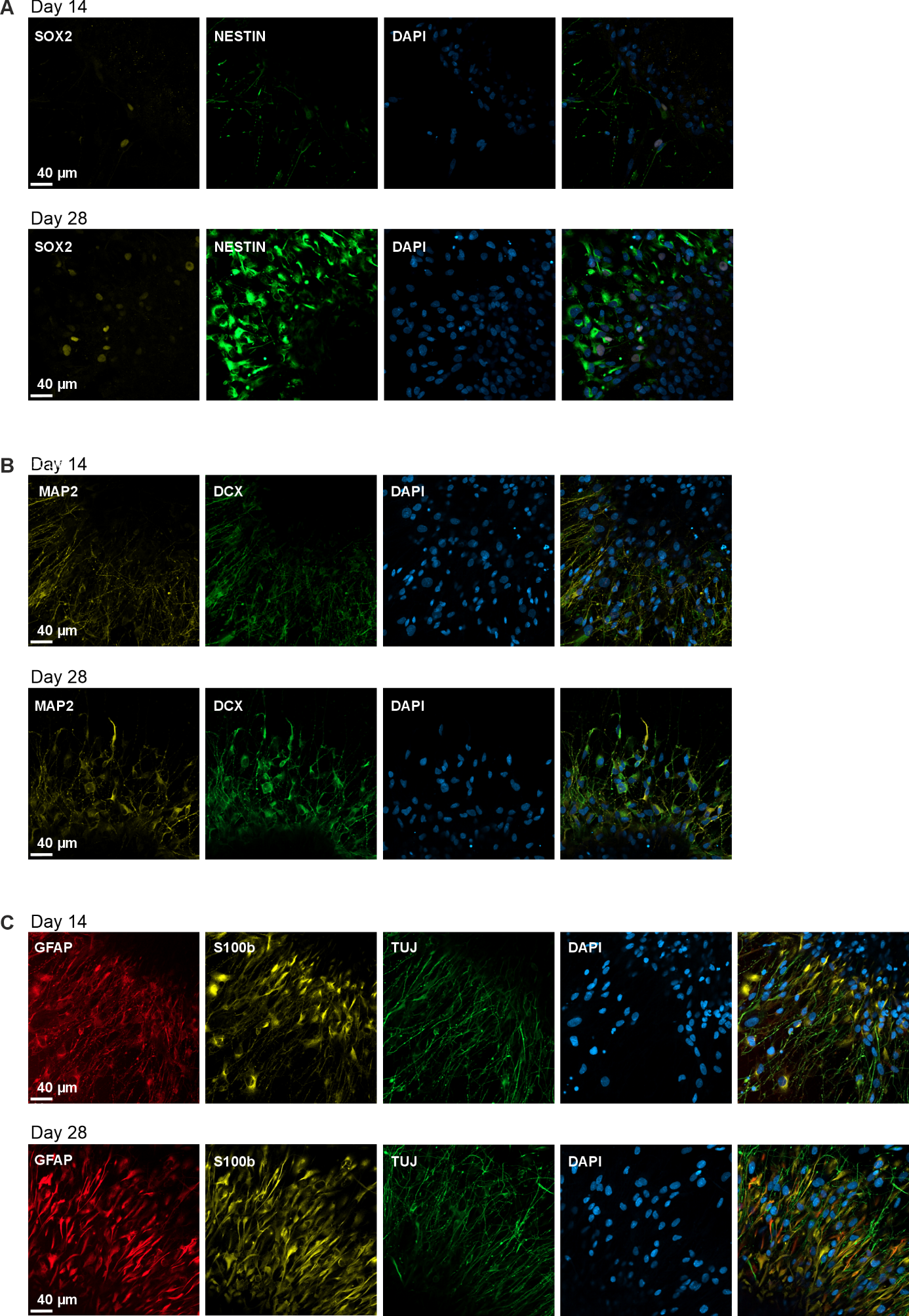

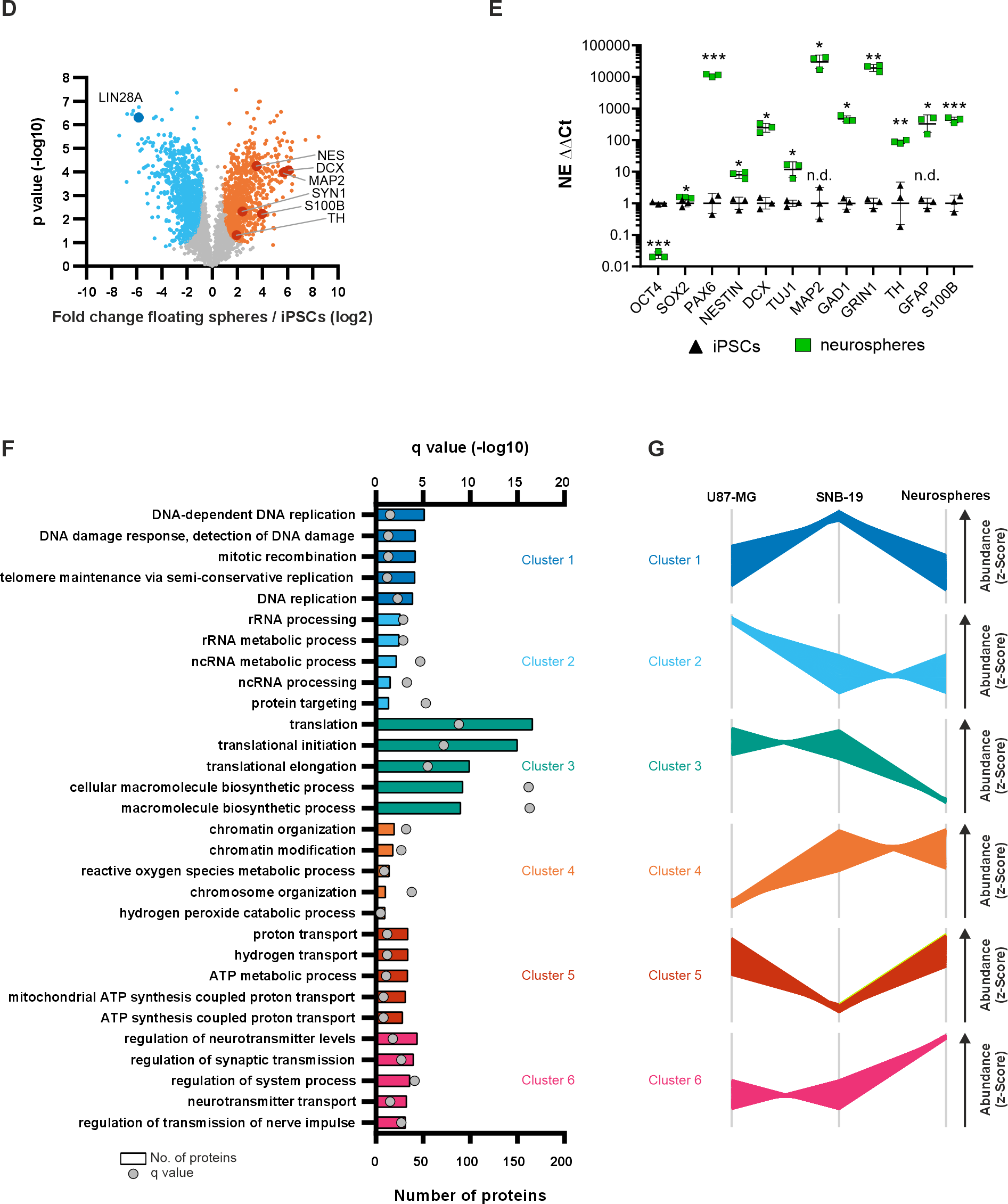

**(A-C)** Confocal immunofluorescence microscopy of 14 or 28 days old spheres stained for (A) SRY-Box Transcription Factor 2 (SOX2) and Nestin, (B) Microtubule-Associated Protein 2 (MAP2) and Doublecortin (DCX), (C) Glial Fibrillary Acidic Protein (GFAP), S100 Calcium Binding Protein B (S100b), and Tubulin Beta-3 (TUJ), and DNA (DAPI). Scale bar: 40 µm. (**D**) Volcano plot showing the comparison of floating neurospheres and iPSCs in whole cell proteome by pairwise t‑tests (3,176 proteins). Proteins with significantly altered abundance in neurospheres (FDR ≤ 0.005, S0 = 0.5, 1,618 proteins) are highlighted in blue (reduced) or orange (increased). Differentially expressed stem cell and neural marker are labelled. (**E**) Gene expression levels of the pluripotency marker Octamer-Binding Transcription Factor 4 (*OCT4*), the neural stem cell and neurodevelopmental regulator *SOX2*, and Paired Box Gene 6 (*PAX6*), the neuronal progenitor cell markers *Nestin* and *DCX*, the neuronal markers *MAP2* and Tubulin Beta-3 (*TUJ1*), the glial markers S100B and GFAP, the GABAergic neuron marker Glutamate Decarboxylase 1 (*GAD1*), the glutamatergic synapse marker Glutamate Ionotropic Receptor NMDA Type Subunit 1 (*GRIN1*), and the dopaminergic neuron marker Tyrosine Hydroxylase (*TH*) were assessed in neurospheres in comparison to iPSCs via qPCR analysis. Data represent geometric mean ± geometric SD; n = 3 biological replicates, 3 technical replicates each. Statistical significance was determined by unpaired t-test. * P ≤ 0.05, ** P ≤ 0.01, *** P ≤ 0.001, **** P ≤ 0.0001. n.d., target gene not detected. (**F-G**) Whole cell proteome analyses comparing global protein expression in floating neurospheres, stem cells (iPSCs) and glioblastoma cells (SNB 19, U87 MG) (24 hpi, n = 4). **(F)** Protein functions (GOBP) enriched among proteins differentially expressed in neurospheres and glioblastoma cells (one‑way ANOVA, FDR ≤ 0.05, 2,902 proteins). Six clusters of differentially expressed proteins were defined by hierarchical clustering (see Figure 1F). A Fisher exact test was applied to determine protein functions enriched (q ≤ 0.05) in each cluster relative to all differentially expressed proteins (3,545 proteins). Only most significant results (q) are shown. Bars display the number of proteins with the respective protein function. Q values are displayed by grey dots. **(**G**)** Profile plots showing the protein abundance (z‑transformed LFQ values) of all proteins belonging to clusters 1‑6 shown in Figure 1F.

**Fig. S2:**

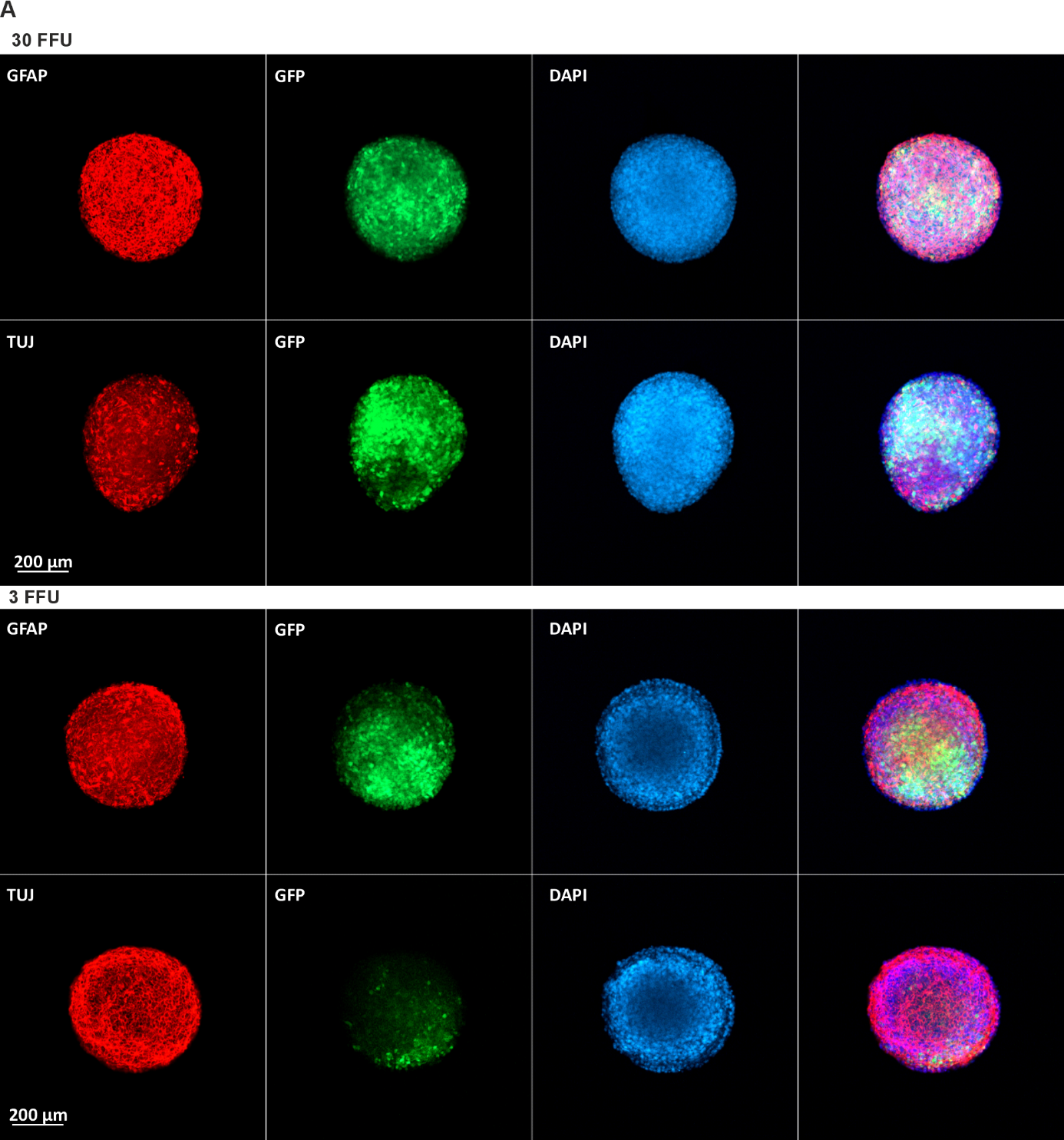

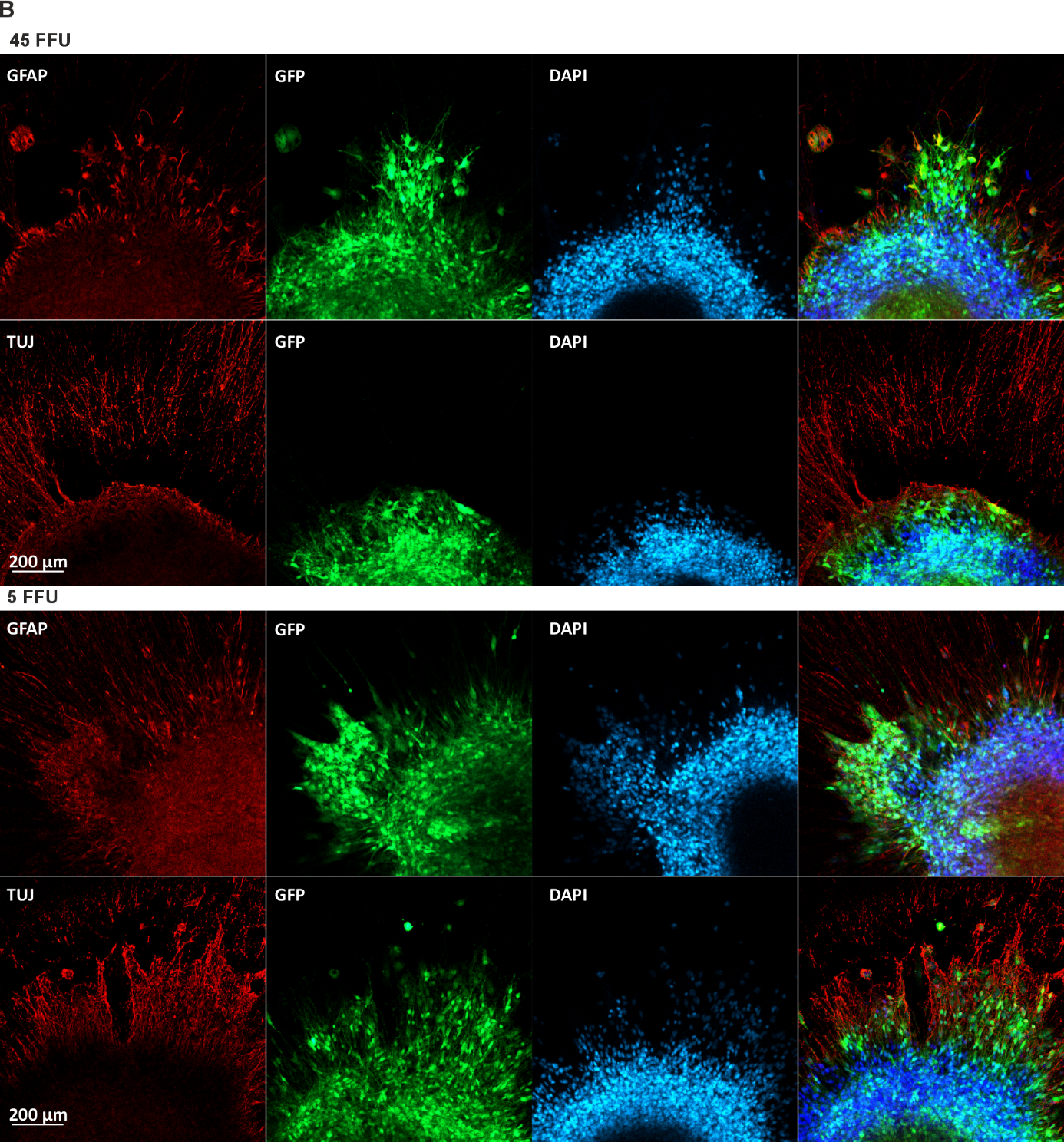

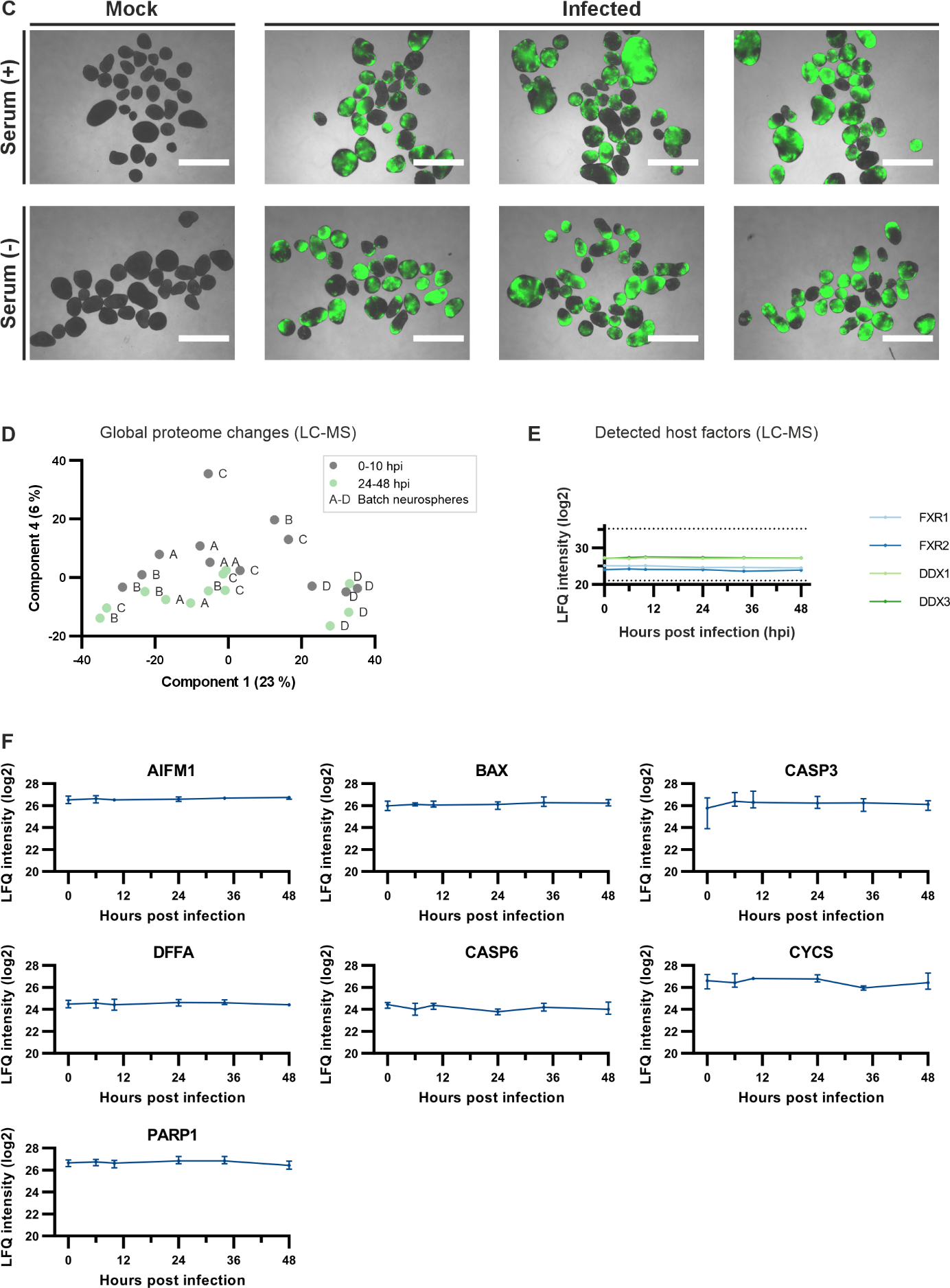

**Infection of floating and plated neurospheres with VEEV.** **(A)** Confocal immunofluorescence microscopy of floating spheres 24 hpi, infected (GFP) with 30 (upper panel) or 3 FFU (lower panel) stained for GFAP, TUJ1 and DNA (DAPI). Scale bar: 200 µm. **(B)** Confocal immunofluorescence microscopy of plated spheres 24 hpi, infected (GFP) with 45 (upper panel) or 5 FFU (lower panel) stained for GFAP, TUJ1 and DNA (DAPI). Scale bar: 200 µm. **(C)** Representative brightfield-fluorescence overlay images of mock- and VEEV-infected neurospheres cultured under serum-containing (Serum (+)) conditions as used for whole cell proteome analyses, and serum-free (Serum (−)) conditions as used for secretome analyses (see Fig. 5). VEEV expression is visualized by GFP expression (green). Images were acquired 24 hpi. Scale bars = 1250 µm. **(D‑E)** Whole cell proteome analysis of floating neurospheres 0‑48 h post VEEV infection (30 FFU per sphere) analyzed by LC‑MS (2,772 proteins). (**D**) Principal component analysis comparing the overall proteomes of floating neurospheres 0‑48 hpi. Letters A‑E indicate the batch of neural progenitor cells the neurospheres were derived from. (**E**) Expression profile of known VEEV host factors over time detected by LC-MS analysis (LFQ intensities). Dotted lines represent minimum and maximum values detected in the selected dataset. (**F**) LFQ intensities of filtered apoptosis markers and PARP1, a DNA repair enzyme recently shown to contribute to motor neuron death^1^, as detected in the whole cell proteome of neurospheres 0-48 hours post VEEV infection (LC-MS, n = 4, non-imputed data).

**Fig. S3:**

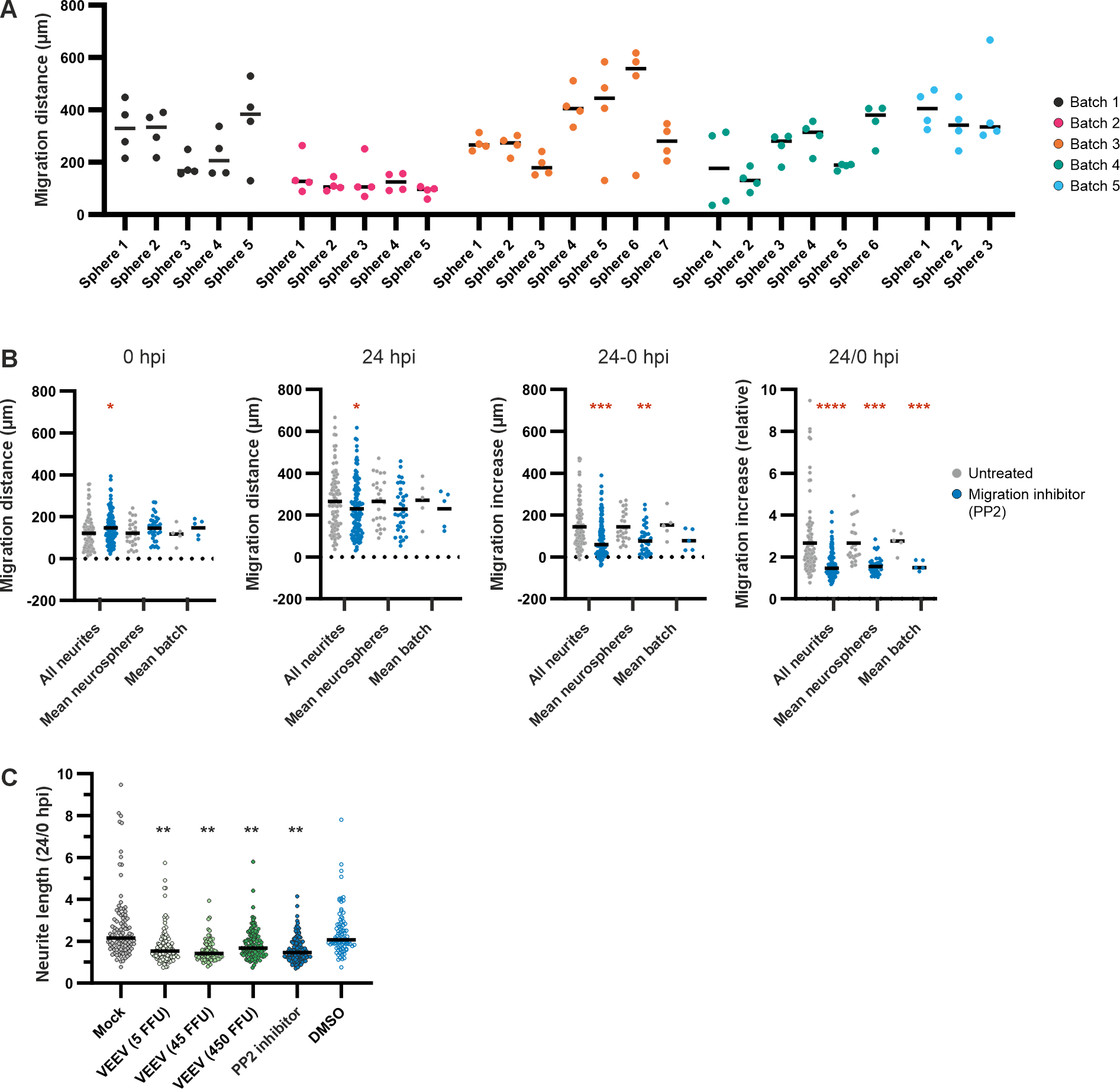

Neurite outgrowth of single plated neurospheres infected by 5, 45, or 450 infectious viral particles (FFU) per sphere. A migration inhibitor (PP2), DMSO and uninfected cells (Mock) served as controls. **(A)** Length of all neurites measured in untreated neurospheres 48 h post plating showing variance of neurite length within spheres, across spheres and across separate experiments resulting from different batches of neural progenitor cells. **(B)** Data distribution of neurite lengths derived from neurospheres treated with and without PP2‑inhibitor. Individual experiments (n = 5) were derived from separate batches of neural progenitor cells. Data distribution is shown for neurite lengths before treatment (0 hpi), 24 h after treatment (24 hpi) and neurite lengths 24 hpi normalized to lengths 0 hpi (24-0 hpi, 24/0 hpi). Each graph shows the data distribution of neurite lengths measured across all experiments (all neurites), of neurite lengths averaged per neurite (mean neurosphere) or averaged per individual experiment (mean batch). Untreated samples were compared to treated samples by pairwise t‑tests. Displayed medians are derived from 104 untreated spheres and 131 inhibitor‑treated spheres across all experiments. **(C)** Lengths of all measured neurites across all experiments as used for calculation of means as displayed in Figure 3D. Neurite outgrowth of untreated spheres (Mock) compared to VEEV‑infected, migration inhibitor‑treated (PP2), or DMSO‑treated (inhibitor solvent) spheres. Neurite lengths measured 24 h post treatment were normalized to neurite lengths measured before treatment (24/0 hpi). Each condition comprises data from five individual experiments derived from separate batches of neural progenitor cells (40-143 measured neurites per group). Conditions were compared to Mock condition by one‑way ANOVA. **(Statistics)** Dashes in dot plots show medians. Asterisks indicate significance levels of 0.05 (*), 0.005 (**), 10^‑5^ (***) and 10^‑10^ (****).

**Fig. S4:**
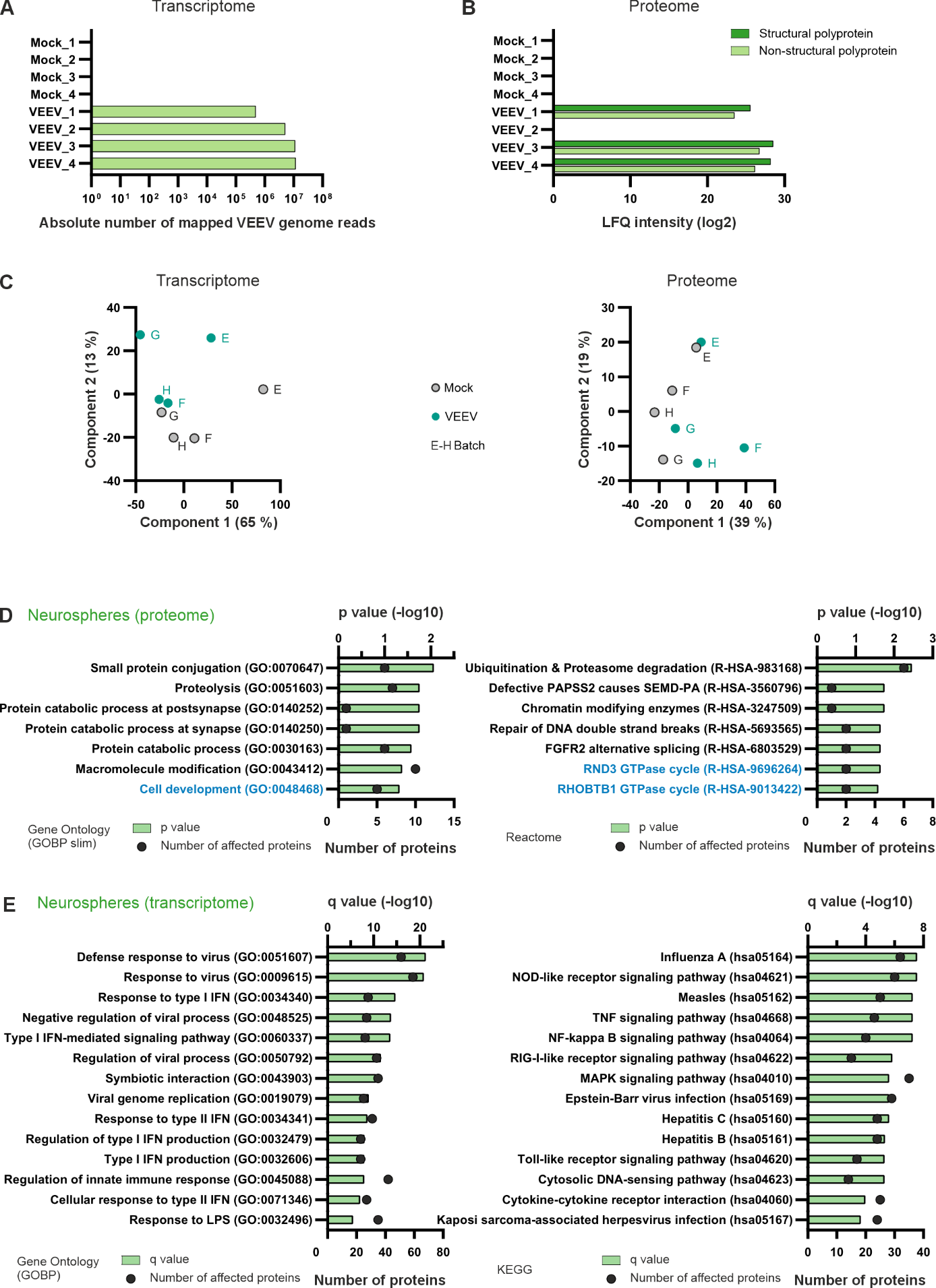

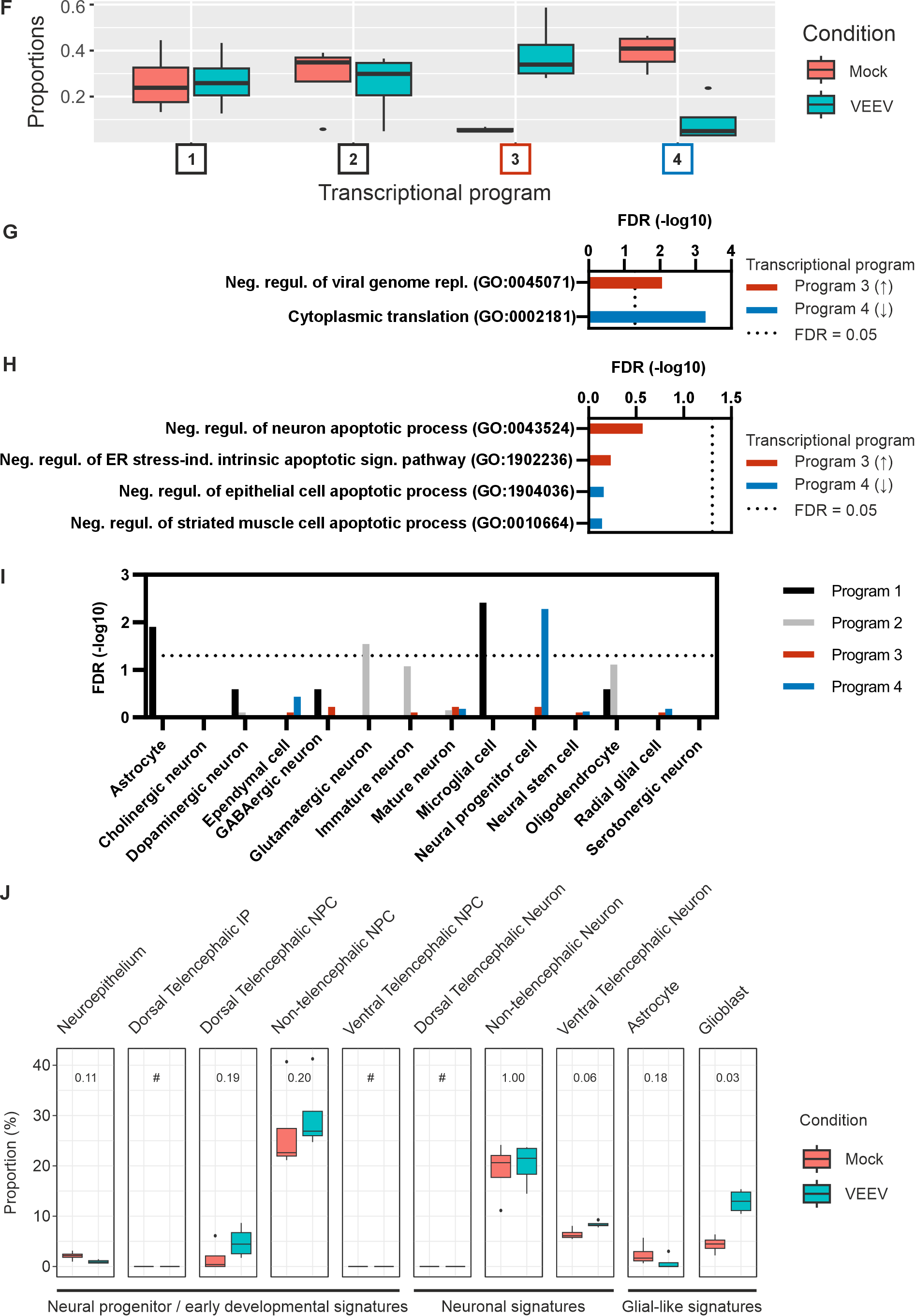

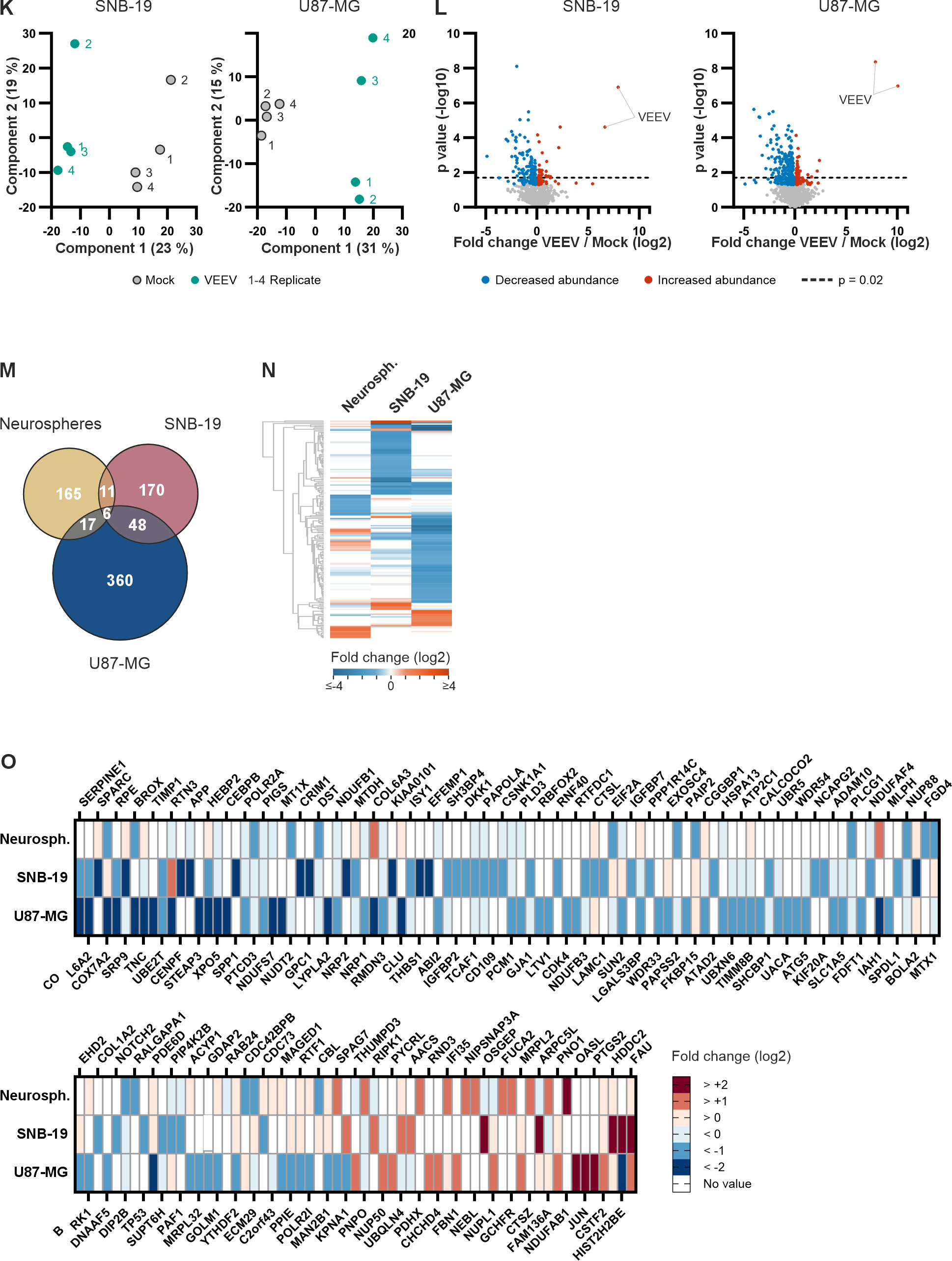

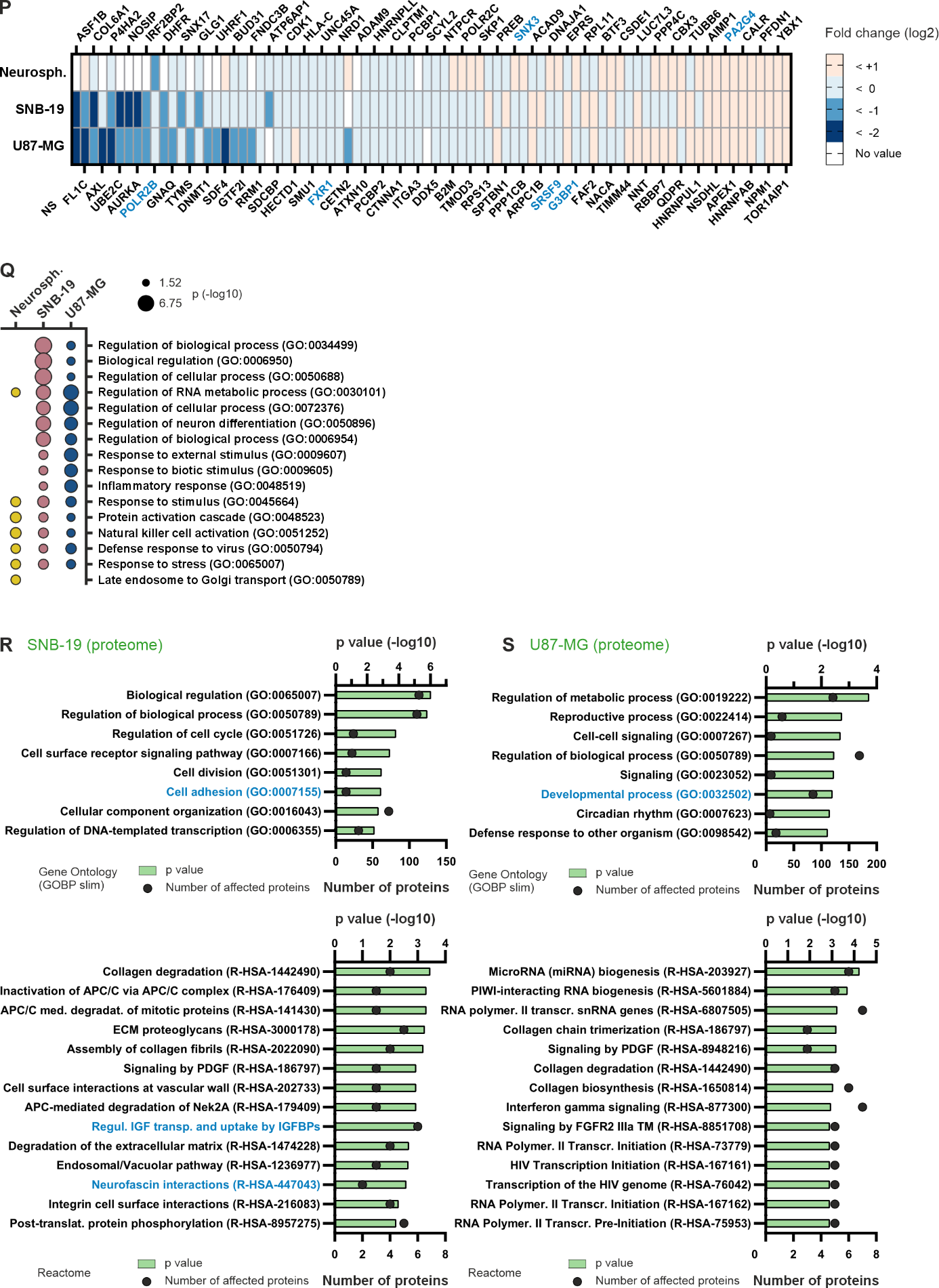

Transcriptome (RNAseq) and whole cell proteome analyses (LC‑MS) of floating neurospheres, SNB‑19 and U87‑MG cells upon VEEV infection (24 hpi, n = 4, 30 FFU per sphere) compared to uninfected cells (Mock). Transcripts and proteins with significantly altered abundance in infected cells (p ≤ 0.05, pairwise t‑tests) were defined as differentially expressed. VEEV proteins were excluded prior to further analyses. Results for glioblastoma cells were derived from experiments under low cell density (experiment 1, compare to secretome analyses, Figure 5). **(A‑B)** Abundance of VEEV transcripts (A, reads mapped to the VEEV genome) and proteins (B, LFQ intensities) in infected and uninfected neurospheres. Different samples were processed for RNAseq‑ and LC‑MS analyses. **(C)** Principal component analysis comparing VEEV‑infected (turquoise) and uninfected (grey) samples analyzed by RNAseq (left) or LC‑MS (right). Letters indicate the batch of neural progenitor cells the neurospheres were derived from. (**D–E**) Cellular processes enriched among differentially expressed proteins (D, p ≤ 0.02) and transcripts (E, p ≤ 0.05) in neurospheres, identified by Fisher exact tests (D: p ≤ 0.01, E: q ≤ 0.001, enrichment factor > 1). Only the most significant terms are shown (sorted by significance). The respective database is indicated at the bottom of each plot. Similar GOBP terms in (E) were grouped using the Revigo algorithm with medium semantic similarity settings. Selected processes are highlighted in blue. (**F**) Unsupervised deconvolution grouped co-regulated genes into shared expression patterns across all replicates and reveals infection-associated shifts in distinct transcriptional programs. Boxplots show the relative contribution (proportions) of four latent gene-expression programs (programs 1-4) across mock (red) and VEEV-infected (blue) neurospheres. (**G-H**) Bar plots showing pathway enrichment (-log10 FDR) for VEEV-responsive DEGs assigned to each transcriptional program following unsupervised deconvolution. (G) displays overall pathway enrichment, while (H) focuses on cell death–related pathways, with the two most significant terms shown per program. The dashed line indicates the significance threshold (FDR = 0.05). (**I**) Bar plots showing cell type enrichment (-log10 FDR) assigned to each transcriptional program following unsupervised deconvolution. (**J**) Boxplots showing shifts of cell-type–associated transcriptional programs upon VEEV infection as estimated by supervised deconvolution. p values (wilcoxon-test) are plotted above the boxes. Supervised deconvolution was performed on RNA-seq data (fpkm) using the Deconomix algorithm trained on the transcriptomic cell atlas of human neural organoids. IP = intermediate progenitor, NPC = neural progenitor cell. **(K)** Principal component analysis comparing VEEV‑infected (turquoise) and uninfected (grey) SNB-19 (left) and U87-MG cells (right) analyzed by LC‑MS. Numbers indicate the biological replicate of the experiment. **(L)** Volcano plots showing overall whole cell proteome changes in SNB‑19 (2,769 proteins) and U87‑MG cells (2,931 proteins). Differentially expressed proteins (201, 237 and 433 proteins) are highlighted in blue (reduced expression) and red (increased expression). Identified VEEV proteins (structural and non‑structural) are labelled. Proteins above the dotted grey line (p ≤ 0.02, 51, 139 and 252 proteins) were used for downstream enrichment analysis shown in R-S. **(M)** Venn diagram showing number of differentially expressed (p ≤ 0.05) proteins in neurospheres, SNB‑19 or U87‑MG cells. **(N)** Heat map showing fold changes of differentially expressed proteins with absolute fold change ≥ 2 (neurospheres, SNB‑19 or U87‑MG cells, 177 proteins). Proteins were grouped by hierarchical clustering with missing values replaced by zero. **(O)** Differentially expressed proteins with absolute fold changes ≥ 2 in neurospheres, SNB‑19 or U87‑MG cells (155 proteins, as shown in heat map in Figure 4B). Fold changes induced by infection are reflected by colors (blue = decreased, red = increased expression). **(P)** Proteins differentially expressed in more than one model (82 proteins, compare to Venn diagram shown in (M). Fold changes induced by infection are reflected by colors (blue = decreased, red = increased expression). Proteins differentially expressed in all three models are highlighted in blue. Proteins are ordered by the sum of fold changes. **(Q)** Comparison of biological processes enriched among proteins differentially expressed (p ≤ 0.05, compare to Venn diagram shown in (M)) in neurospheres, SNB‑19 or U87‑cells (GOBP, Fisher exact test, 6 most significant enriched processes of each cellular model). Similar GOBP entries were grouped using Revigo (slim semantic similarity settings). Significance levels are reflected by the size of bubbles. (**R-S**) Cellular processes enriched (Fisher exact test, p ≤ 0.01, enrichment factor > 1) among proteins differentially expressed (p ≤ 0.02, as shown in L) in SNB‑19 cells (R) or U87‑MG cells (S). Only most significant results are shown (sorted by p value). The respective database is indicated at the bottom of the plot. Similar GOBP processes were grouped using Revigo (medium semantic similarity settings). Selected processes are highlighted in blue.

**Fig. S5:**

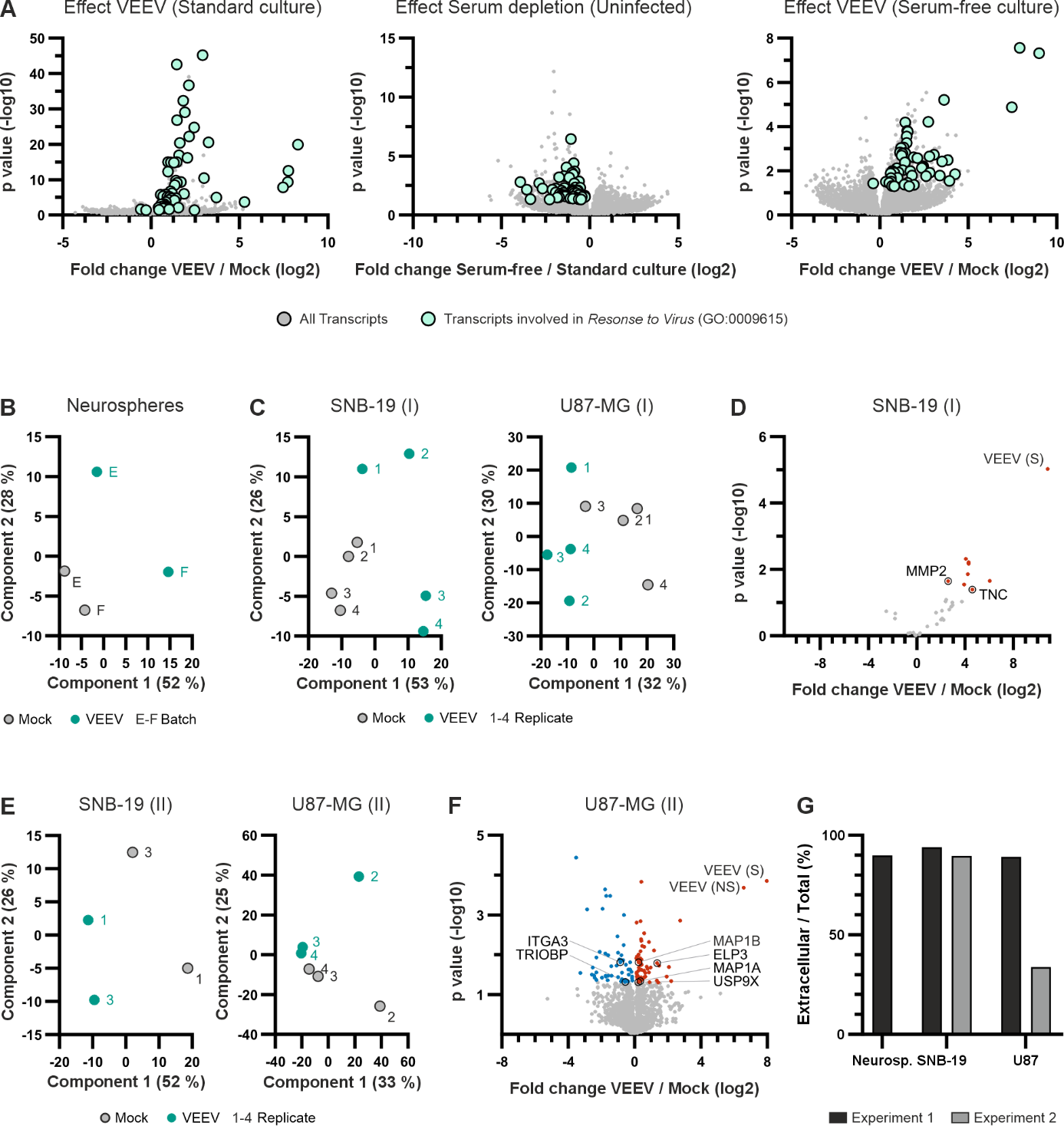

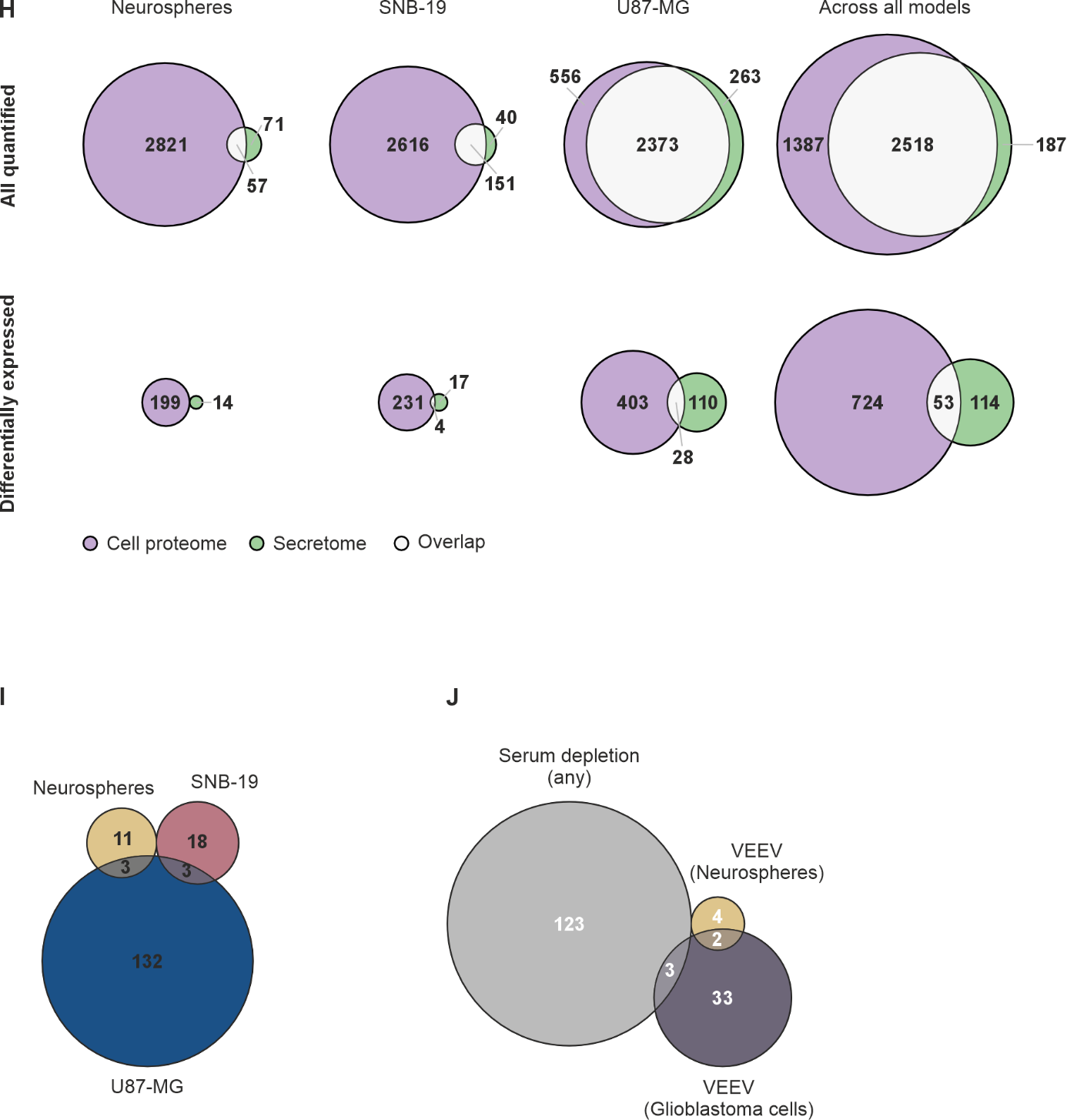

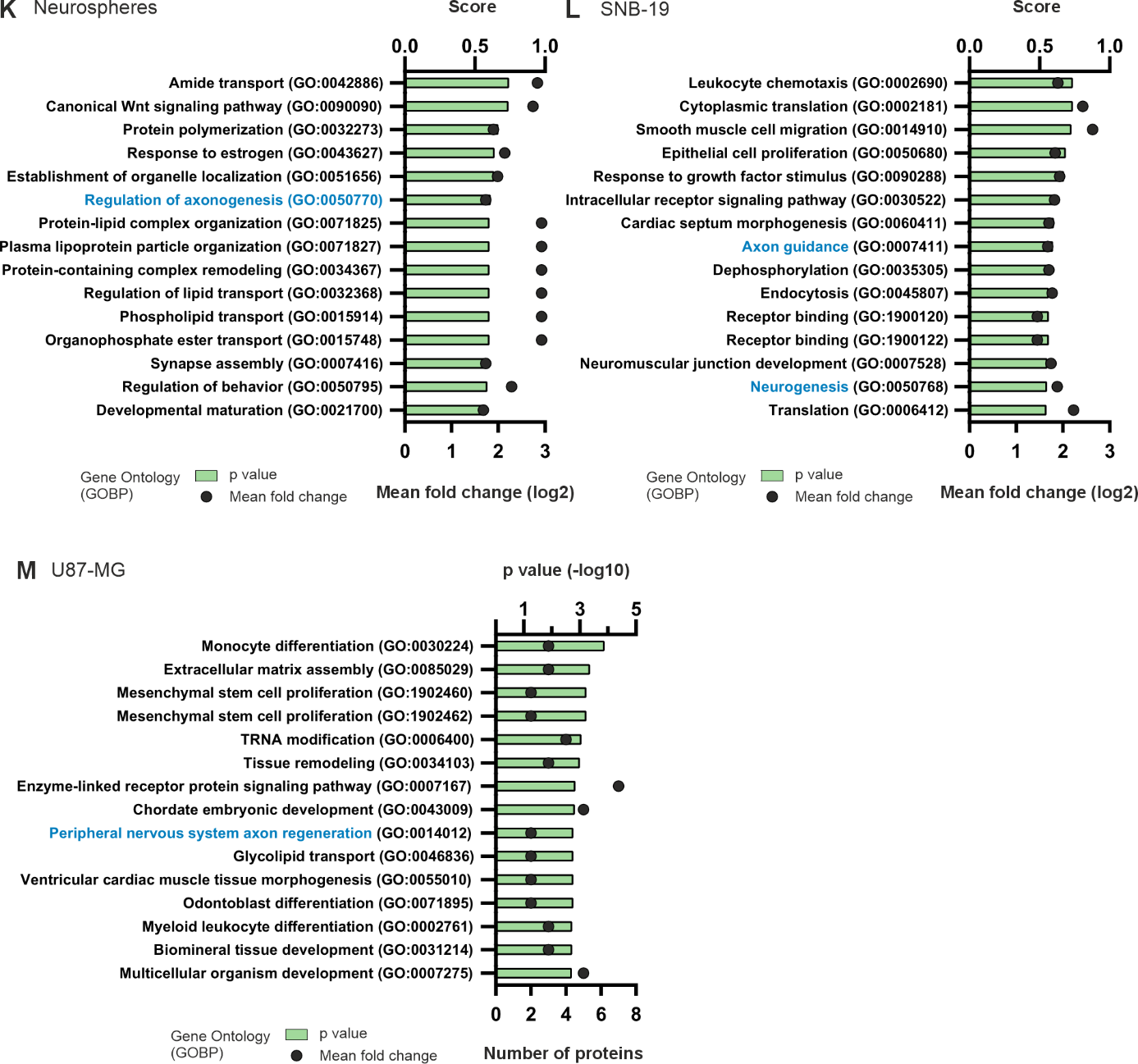

Secretome analyses of floating neurospheres, SNB‑19 and U87‑MG cells upon VEEV infection (24 hpi, n = 4, 30 FFU per sphere) under serum‑free conditions compared to uninfected cells (Mock). A single experiment was performed with neurospheres (n = 2 after quality control). Experiments with glioblastoma cells were performed under low (experiment 1, n = 4) and high cell density condition (experiment 2, n = 2 for SNB‑19, n = 3 for U87‑MG). Proteins with significantly altered abundance in infected cells (p ≤ 0.05, pairwise t‑tests) and proteins with absolute fold changes ≥ 2 in both biological replicates (neurospheres only) were defined as differentially secreted. Results of glioblastoma experiments were grouped. Effect of serum depletion was estimated based on transcriptome changes induced by serum‑depletion. **(A)** Comparison of transcriptome changes induced by VEEV infection in neurospheres (standard culture conditions or serum‑free, compare to Figure 4) to changes induced by serum depletion. Volcano plots show overall transcriptome changes (35,905 35,968 and 35,493 transcripts). Differentially expressed transcripts (p ≤ 0.05) involved in defense mechanisms against viruses (Gene Ontology GO:0009615) are highlighted in turquoise (91 transcripts). n = 4, except for Mock serum‑free (n = 3). **(B)** Principal component analysis comparing the overall secretome of VEEV‑infected (turquoise) and uninfected (grey) neurospheres analyzed by LC‑MS. Letters indicate the batch of neural progenitor cells the neurospheres were derived from. **(C)** Principal component analysis comparing the overall secretomes (experiment 1) of VEEV‑infected (turquoise) and uninfected (grey) SNB-19 (left) and U87-MG cells (right) analyzed by LC‑MS. Numbers indicate the biological replicate of the experiment. **(D)** Volcano plot showing changes in the secretome of SNB-19 cells (experiment 1, n = 4, 33 proteins). Differentially secreted proteins (9) are highlighted in blue (reduced) or red (increased). Differentially secreted proteins annotated as modulators of neural cell migration or neurite outgrowth are highlighted by black circles and labelled. Proteins significantly reduced in neurospheres and U87-MG cells are labelled in bold letters. Viral proteins (structural (S) and non-structural (NS) proteins) are labelled in grey letters. **(E)** Principal component analysis comparing the overall secretomes (experiment 2) of VEEV‑infected (turquoise) and uninfected (grey) SNB-19 (left) and U87-MG cells (right) analyzed by LC‑MS. Numbers indicate the biological replicate of the experiment. **(F)** Volcano plot showing changes in the secretome of U87-MG cells (experiment 2, n = 3, 2,568 proteins). Differentially secreted proteins (116) are highlighted in blue (reduced) or red (increased). Differentially secreted proteins annotated as modulators of neural cell migration or neurite outgrowth are highlighted by black circles and labelled. Proteins significantly reduced in neurospheres and U87-MG cells are labelled in bold letters. Viral proteins (structural (S) and non-structural (NS) proteins) are labelled in grey letters. **(G)** Proportion of all analyzed secretome proteins in different experiments that are annotated as extracellular (GOCC slim database). Different experiments performed are represented by black and grey bars. **(H)** Venn diagrams showing the number of proteins detected (upper panel) or differentially expressed (lower panel) in the cell proteome (purple), the secretome (green) or in both (white). For glioblastoma cells, results of experiments 1 and 2 were grouped. For the right panel (across all models), proteins derived from experiments with neurospheres, SNB‑19 and U87‑MG cells were grouped. **(I)** Venn diagram showing the number of differentially secreted protein by neurospheres, SNB‑19 or U87‑MG cells. **(J)** Comparison of secretome changes induced by VEEV infection in neurospheres or glioblastoma cells to whole cell proteome changes induced by serum depletion (any experiment). Whole cell proteomes under serum‑free conditions were derived from secretome experiments and were compared to whole proteomes under serum‑containing conditions derived from whole cell proteome experiments shown in Figure 4 (uninfected samples). Venn diagram is showing the numbers of proteins with significantly altered abundance (p 0.05, absolute fold change ≥ 2). Proteins affected by serum depletion in neurospheres (1 experiment), SNB‑19 cells (2 experiments) or U87‑MG cells (2 experiments) were grouped, as well as proteins affected by VEEV in SNB‑19 cells or U87‑MG cells (each 2 experiments). **(K-M)** Affected cellular processes (GOBP database) in neurospheres (K), SNB‑19 cells (L) and U87‑MG cells (M). (K-L) Enrichment among proteins with VEEV‑induced absolute fold changes above average ranked by Score (1D enrichment). (M) Enrichment among differentially secreted proteins (p ≤ 0.02) ranked by p value (Fisher exact test). Similar processes were grouped (medium semantic similarity settings). Only 15 most significant enriched processes are displayed. Processes potentially related to neural cell migration or neurite outgrowth are highlighted in blue.

**Fig. S6:**

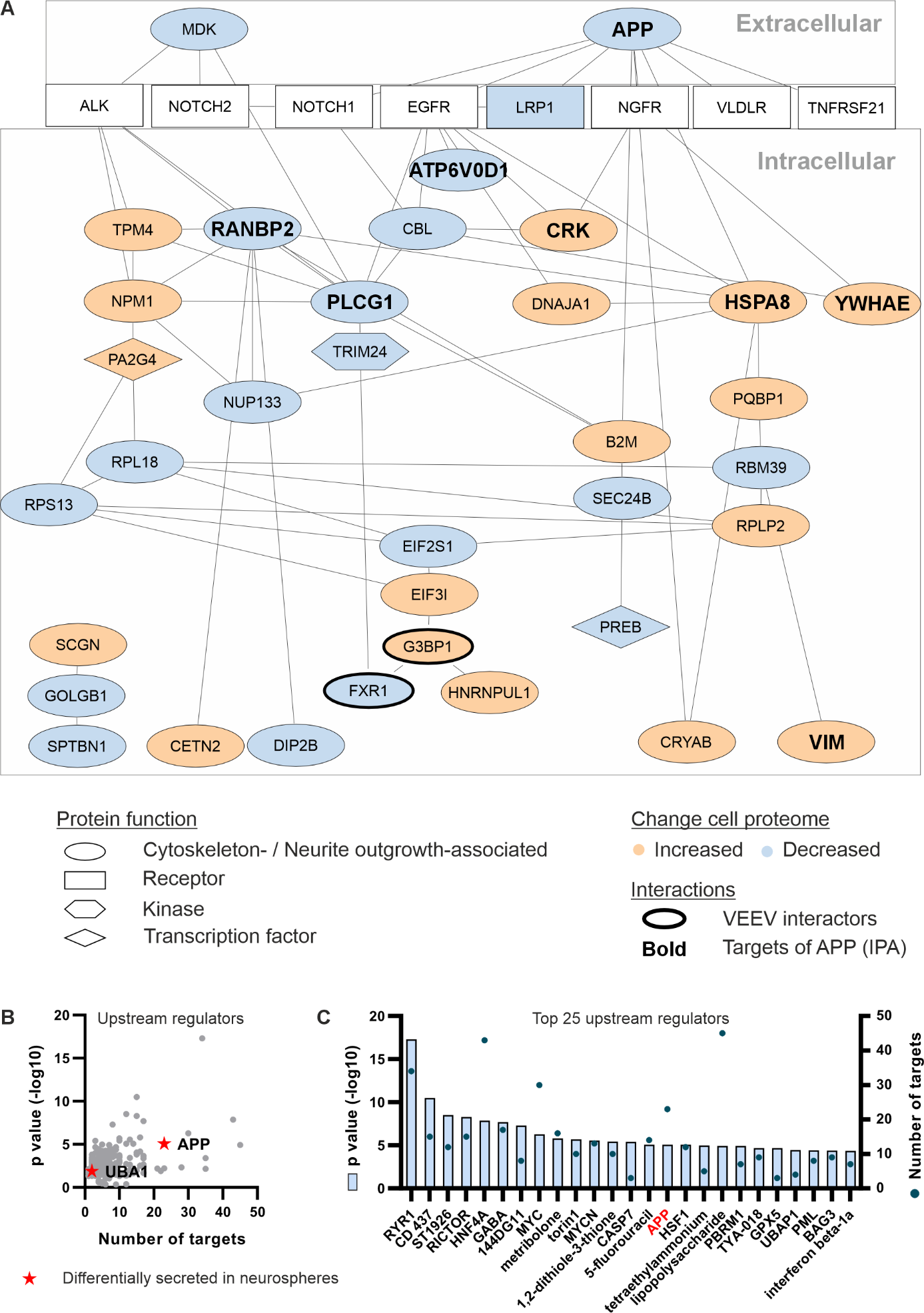

(**A**) Network linking secreted ligands to intracellular effectors of cytoskeleton remodeling and neurite outgrowth, kinases and transcription factors that are differentially expressed in VEEV-infected neurospheres (30 FFU, 24 hpi, Table S9-10). The protein–protein interaction network shows connections between differentially secreted ligands (MDK, APP), their candidate cell-surface receptors (any, as detetcted by RNAseq), and downstream intracellular proteins identified in the whole-cell proteome. Nodes represent proteins and edges indicate known interactions (STRING database), highlighting signaling routes converging on cytoskeleton- and neurite-related effectors. The upstream regulator APP and downstream targets as identified by unbiased analysis (B-C) are highlighted by bold and bigger letters. (**B**) Results of upstream regulator analysis with Ingenuity Pathway Analysis software (IPA, QIAGEN). The list of intracellular proteins differentially expressed upon VEEV infection (199) served as input data, threshold was set to p = 0.05. The plot shows probabilities and number of downstream targets of all identified upstream regulators. Proteins identified to be differentially secreted upon VEEV infection are highlighted by red stars. (**C**) Top 25 upstream regulators (IPA).

**Supplementary Tables**

Table S1: neural induction medium (NIM)

| **Component** | **Concentration** |
| --- | --- |
| DMEM-medium, high glucose, Thermo Fisher Scientific #31966-021 | 2 parts |
| Ham’s F12-medium, Thermo Fisher Scientific #31765-027 | 1 part |
| Knockout Serum Replacement (KSR), Thermo Fisher Scientific #10828-028 | 20% |
| N21 supplement without Vitamin A, Miltenyi #130-093-263 | 2% |
| N2 supplement, Thermo Fisher Scientific #17502-048 | 1% |
| Penicillin-Streptomycin (10.000 U/mL), Thermo Fisher Scientific #15140122 | 1% |
| hEGF, Thermo Fisher Scientific #PHG0313 | 20 ng/mL |
| Additives D0-D11: |  |
| SB-431542 hydrate, MedChemExpress/Hycultec #S4317 | 10 μM |
| LDN-193189 hydrochloride, Sigma Aldrich #SML0559 | 0.5 μM |
| Additives D12-D21: |  |
| bFGF, R&D #233-FB | 20 ng/mL |
| Additives during seeding of cells |  |
| Y-27632 TargetMol #T1725 | 10 µM |

Table S2: CINDA+ medium

| **Component** | **Concentration** |
| --- | --- |
| DMEM-medium, high glucose, Thermo Fisher Scientific #31966-021 | 2 parts |
| Ham’s F12-medium, Thermo Fisher Scientific #31765-027 | 1 part |
| Penicillin-Streptomycin (10.000 U/mL), Thermo Fisher Scientific #15140122 | 1% |
| B27 Plus Supplement, Thermo Fisher Scientific #A35828-01 | 2% |
| N2 Supplement Thermo Fisher Scientific #17502-048 | 1% |
| hEGF, Thermo Fisher Scientific #PHG0313 | 20 ng/mL |
| Recombinant human interferon γ, Peprotech #300-02 | 0.05 mg/mL |
| Recombinant human neutropin, Peprotech #450-03 | 20 µg/mL |
| l-ascorbic acid, Sigma Aldrich #A5960 | 150 mM |
| Creatine monohydrate, Sigma Aldrich #C3630 | 650 mg/mL |
| Freshly added (stable up to 7 days): |  |
| Dibutyryl CyclicAMP sodium salt, Sigma Aldrich #D0260 | 300 µM |

Table S3: Antibody dilutions

| **Target/ Description** | **Dilution** | **Source** |
| --- | --- | --- |
| GFAP, rabbit | 1:200 | Atlas antibodies #HPA056030 |
| GFAP, chicken | 1:1000 | Abcam  #ab4674 |
| TUJ, rabbit | 1:100 | Sigma Aldrich #T2200XX |
| DCX, mouse | 1:200 | Thermo Fisher  #MA5-17066 |
| MAP2, rabbit | 1:50 | Cell Signaling  #4542 |
| S100b, rabbit | 1:200 | Thermo Fisher  # MA5-42438 |
| Nestin, mouse | 1:200 | R&D  # MAB1259 |
| Sox2, rabbit | 1:200 | Cell Signaling  #3579 |
| Alexa Fluor™ 568, rabbit | 1:1000 | Invitrogen #A11036 |
| Alexa Fluor™ 647, chicken | 1:1000 | ThermoFisher #A21449 |
| Alexa Fluor™ 488, mouse | 1:1000 | Invitrogen  #A11001 |

Table S4: image processing

| **Image ID; original data** | **shown in figure** | **channel** | **Min** | **Max** |
| --- | --- | --- | --- | --- |
| Migrating Spheres | | | | |
| 20230922_KC_GFAP-MOCK_24hpi_B4.czi | Fig 1 | AF 568 | 4 | 110 |
|  |  | DAPI | 2 | 116 |
| 20230922_KC_TUJ1-MOCK_24hpi_B2.czi | Fig 1 | AF 568 | 3 | 126 |
|  |  | DAPI | 2 | 95 |
| 20230630_V1_SF13_VEEV_24hpi_E4_10-3_GFAP.czi | Fig 2 | AF 568 | 1 | 41 |
|  |  | GFP | 1 | 106 |
|  |  | DAPI | 1 | 135 |
| 20230921_KC_TUJ1-VEEV 10^-3_E3.czi | Fig 2 | AF 568 | x | x |
| No adjustment |  | GFP | x | x |
|  |  | DAPI | x | x |
| Day14_Nestin_sox2_I | SI Fig 1 | AF 568  AF 488  DAPI | 0  2  4 | 161  91  140 |
| Day28_Nestin_Sox2_II | SI Fig 1 | AF 568  AF 488  DAPI | 0  2  4 | 161  91  140 |
| Day14_DCX_MAP2_I | SI Fig 1 | AF 568  AF 488  DAPI | 0  10  0 | 190  190  201 |
| Day28_DCX_MAP2_I | SI Fig 1 | AF 568  AF 488  DAPI | 0  10  0 | 190  190  201 |
| Day14_TUJ_GFAP_s100b_I | SI Fig 1 | AF 647  AF 568  AF 488  DAPI | 0  2  0  0 | 104  91  135  238 |
| Day28_TUJ_GFAP_s100b_I | SI Fig 1 | AF 647  AF 568  AF 488  DAPI | 0  2  0  0 | 104  91  135  104 |
| 20230630_V1_SF13_VEEV_24hpi_D3_10-4_GFAP.czi | SI Fig 2 | AF 568 | 1 | 54 |
|  |  | GFP | 1 | 54 |
|  |  | DAPI | 1 | 97 |
| 20230630_V1_SF13_VEEV_24hpi_D4_10-4_TUJ1.czi | SI Fig 2 | AF 568 | 1 | 93 |
|  |  | GFP | 1 | 68 |
|  |  | DAPI | 1 | 138 |
| 20230630_V1_SF13_VEEV_24hpi_C3_10-5_GFAP_02.czi | SI Fig 2 | AF 568 | 1 | 42 |
|  |  | GFP | 1 | 102 |
|  |  | DAPI | 1 | 87 |
| 20230630_V1_SF13_VEEV_24hpi_C5_10-5_TUJ1.czi | SI Fig 2 | AF 568 | 8 | 68 |
|  |  | GFP | 2 | 48 |
|  |  | DAPI | 1 | 101 |
| Floating spheres | | | | |
| KC_GFAP-MOCK_24hpi_whole-sphere.czi | SI Fig 1 | AF568 | 15 | 63 |
|  |  | DAPI | 13 | 97 |
| KC_TUJ1-MOCK_24hpi_whole-sphere.czi | SI Fig 1 | AF 568 | 18 | 169 |
|  |  | DAPI | 13 | 114 |
| KC_GFAP-VEEV 10^-3 24hpi_whole-sphere.czi | SI Fig 2 | AF 568 | 1 | 57 |
|  |  | GFP | 0 | 255 |
|  |  | DAPI | 9 | 123 |
| KC_TUJ1-VEEV 10^-3 24hpi_whole-sphere.czi | SI Fig 2 | AF 568 | 1 | 93 |
|  |  | GFP | 10 | 201 |
|  |  | DAPI | 9 | 121 |
| KC_GFAP-VEEV 10^-4 24hpi_whole-sphere.czi | SI Fig 2 | AF 568 | 52 | 255 |
|  |  | GFP | 6 | 210 |
|  |  | DAPI | 10 | 159 |
| KC_TUJ1-VEEV 10^-4 24hpi_whole-sphere.czi | SI Fig 2 | AF 568 | 16 | 103 |
|  |  | GFP | 41 | 255 |
|  |  | DAPI | 10 | 155 |
| KC_GFAP-VEEV 10^-5 24hpi_whole-sphere.czi | SI Fig 2 | AF 568 | 22 | 99 |
|  |  | GFP | 7 | 75 |
|  |  | DAPI | 9 | 108 |
| KC_TUJ1-VEEV 10^-5 24hpi_whole-sphere.czi | SI Fig 2 | AF 568 | 11 | 128 |
|  |  | GFP | 10 | 53 |
|  |  | DAPI | 11 | 123 |

Table S5: RNA concentration after RNA preparation

| **Batch** | **VEEV infection (hpi)** | **RNA conc. (ng/µL)** | **Total RNA per sample (ng)** |
| --- | --- | --- | --- |
| I | 00 hpi | 27.6 | 1104 |
| I | 06 hpi | 18.3 | 732 |
| I | 10 hpi | 17.4 | 696 |
| I | 24 hpi | 40.3 | 1612 |
| I | 34 hpi | 26.2 | 1048 |
| I | 48 hpi | 30.6 | 1224 |
| II | 00 hpi | 17.6 | 704 |
| II | 06 hpi | 2.7 | 108 |
| II | 10 hpi | 5.0 | 200 |
| II | 24 hpi | 9.1 | 364 |
| II | 34 hpi | 5.0 | 200 |
| II | 48 hpi | 3.8 | 152 |
| III | 00 hpi | 16.2 | 648 |
| III | 06 hpi | 12.2 | 488 |
| III | 10 hpi | 17.9 | 716 |
| III | 24 hpi | 28.4 | 1136 |
| III | 34 hpi | 18.3 | 732 |
| III | 48 hpi | 11.5 | 460 |

Table S6: RT‑qPCR Mix

| **Component** | **Note** | **Volume per reaction (μL)** | **Final concentration** |
| --- | --- | --- | --- |
| RNAse-free water |  | 1.25 |  |
| 2x Reaction mix |  | 6.25 | 1x |
| VEEV_forward: 5`-ACC ATG CTA ATG CCA GAG CGT TTT CGC A-3 | Pre-Mixed | 1.00 | 0.50 μM |
| VEEV reverse: 5´- GGG CGC ACT TCC AAT GTC AAG GAT-3´ |  |  | 0.50 μM |
| VEEV Probe: FAM- TGA TCG AAA CGG AGG TGG ACC CAT CC-TAMRA |  |  | 0.25 μM |
| *ACTB* forward: AGC GCA AGT ACT CCG TGT G | Pre-Mixed | 1.00 | 0.20 μM |
| *ACTB* reverse: CGG ACT CAT CGT ACT CCT GCT T |  |  | 0.20 μM |
| *ACTB* Probe: HEX- TCG CTG TCC ACC TTC CAG CAG ATG T-BHQ1 |  |  | 0.20 μM |
| 25x RT-PCR enzyme mix |  | 0.50 | 1x |
| Template RNA |  | 2.50 | 6.25 % of total purified RNA per PCR reaction (neurospheres’ samples)  or  1 × 10^1^ to 1 × 10^8^ genome copies per microliter (VEEV standard) |

Table S7: Thermocycler program

| **Step** | **Time** | **Temperature (°C)** | **Cycles** |
| --- | --- | --- | --- |
| RT | 10 min | 45 | 1 |
| Activation Tag | 10 min | 95 | 1 |
| Denaturation | 15 s | 95 | 42 |
| Annealing | 20 s | 57 |  |
| Elongation | 30 s | 72 |  |
| Storage | ∞ | 20 | 1 |

Table S8: Primer pairs

| **Gene** | **Forward primer (5’-3’)** | **Reverse primer (5’-3’)** | **Product size [bp]** |
| --- | --- | --- | --- |
| OCT4 NM_001173531.3 | CGAGAAGGATGTGGTCCGAG | GGGAAAGGGACCGAGGAGTA | 213 |
| SOX2 NM_003106.3 | TACAGCATGTCCTACTCGCAG | GAGGAAGAGGTAACCACAGGG | 215 |
| PAX6 NM_000280.6 | CCAGCCAGACCTCCTCATAC | TGGCTGACTGTTCATGTGTGTC | 113 |
| NESTIN NM_006617.1 | GCGGGCTACTGAAAAGTTCC | CAGGAGGGTCCTGTACGTGG | 157 |
| DCX NM_000555.3 | GGGAAACCCATCAGCCACAG | TGGAGGTTCCGTTTGCTGAGTC | 122 |
| TUJ1 NM_006086.3 | CTCGAGCCATTCTGGTGGAC | CCCCACTCTGACCAAAGATGA | 108 |
| MAP2 NM_002374.4 | GCTCCCGGAGAAGGATTCTG | CAAGCTGAAGAATCAGCGCA | 82 |
| GAD1 NM_000817.3 | GACCCCAATACCACTAACCTG | GCTGTTGGTCCTTTGCAAG | 117 |
| GRIN1 NM_000832.7 | CTCCTGGAAGATTCAGCTCAA | GTGGATGGCTAACTAGGATGG | 125 |
| TH NM_000360.4 | TGCTAAACCTGCTCTTCTCC | GGTCTCTAGATGGTGGATTTTGG | 104 |
| GFAP NM_002055.5 | CACTGTGAGGCAGAAGCTC | CCTCCAGCGACTCAATCTTC | 137 |
| S100B NM_006272.3 | CACAAGCTGAAGAAATCCGAAC | CACATTCGCCGTCTCCATC | 133 |

Table S9: Annotations associated to cytoskeleton remodeling and neurite outgrowth

| **Database** | **Relevant_terms** |  | **Database** | **Relevant_terms** |
| --- | --- | --- | --- | --- |
| GOBP | actin cytoskeleton organization |  | GOBP | regulation of plasma membrane bounded cell projection organization |
| GOBP | actin filament bundle assembly |  | GOCC | actin cytoskeleton |
| GOBP | actin filament bundle organization |  | GOCC | actin filament |
| GOBP | actin filament capping |  | GOCC | actin filament bundle |
| GOBP | actin filament organization |  | GOCC | actin-based cell projection |
| GOBP | actin filament-based movement |  | GOCC | cell junction |
| GOBP | actin filament-based process |  | GOCC | cell leading edge |
| GOBP | axon guidance |  | GOCC | cell projection |
| GOBP | cell projection assembly |  | GOCC | cell projection membrane |
| GOBP | cell projection organization |  | GOCC | contractile actin filament bundle |
| GOBP | contractile actin filament bundle assembly |  | GOCC | cortical actin cytoskeleton |
| GOBP | cytoskeleton organization |  | GOCC | filopodium |
| GOBP | intermediate filament cytoskeleton organization |  | GOCC | focal adhesion |
| GOBP | microtubule cytoskeleton organization |  | GOCC | glial cell projection |
| GOBP | microtubule cytoskeleton organization involved in mitosis |  | GOCC | growth cone |
| GOBP | negative regulation of actin filament depolymerization |  | GOCC | lamellipodium |
| GOBP | negative regulation of actin filament polymerization |  | GOCC | microtubule cytoskeleton |
| GOBP | negative regulation of cell projection organization |  | GOCC | neuron projection |
| GOBP | negative regulation of cytoskeleton organization |  | GOCC | neuron projection cytoplasm |
| GOBP | negative regulation of neuron projection development |  | GOCC | neuron projection membrane |
| GOBP | neuron projection development |  | GOCC | neuron projection terminus |
| GOBP | neuron projection regeneration |  | GOCC | plasma membrane bounded cell projection |
| GOBP | plasma membrane bounded cell projection assembly |  | GOCC | plasma membrane bounded cell projection cytoplasm |
| GOBP | plasma membrane bounded cell projection organization |  | GOMF | actin filament binding |
| GOBP | positive regulation of cell projection organization |  | GOMF | cytoskeletal motor activity |
| GOBP | positive regulation of neuron projection development |  | GOMF | dynein motor activity |
| GOBP | protein localization to microtubule cytoskeleton |  | GOMF | kinesin motor activity |
| GOBP | regulation of actin cytoskeleton organization |  | GOMF | microtubule motor activity |
| GOBP | regulation of actin filament depolymerization |  | GOMF | myosin motor activity |
| GOBP | regulation of actin filament length |  | GOMF | structural constituent of cytoskeleton |
| GOBP | regulation of actin filament organization |  | KEGG | Axon guidance |
| GOBP | regulation of actin filament polymerization |  | KEGG | Focal adhesion |
| GOBP | regulation of actin filament-based movement |  | KEGG | Regulation of actin cytoskeleton |
| GOBP | regulation of actin filament-based process |  | Keywords | Actin filament bundle |
| GOBP | regulation of cell projection assembly |  | Keywords | Neuron projection |
| GOBP | regulation of cell projection organization |  | Reactome | Axon guidance |
| GOBP | regulation of cytoskeleton organization |  | Reactome | Cell junction organization |
| GOBP | regulation of microtubule cytoskeleton organization |  | Reactome | EPHA-mediated growth cone collapse |
| GOBP | regulation of neuron projection development |  | Reactome | Rho GTPases Activate Formins |
| GOBP | regulation of plasma membrane bounded cell projection assembly |  |  |  |

Table S10: Annotations associated to kinases

| **Database** | **Relevant_terms** |
| --- | --- |
| GOMF | kinase activity |
| GOMF | protein kinase activity |
| GOMF | phosphotransferase activity |
| GOMF | phosphotransferase activity, alcohol group as acceptor |
| GOMF | protein tyrosine kinase activity |
| GOMF | serine/threonine kinase activity |
| Keywords | Kinase |
| Keywords | Protein kinase |
| Pfam | APS_kinase |
| Pfam | Pkinase |
| Pfam | Pkinase_Tyr |
